# Representative diatom and coccolithophore species exhibit divergent responses throughout simulated upwelling cycles

**DOI:** 10.1101/2020.04.30.071480

**Authors:** Robert H. Lampe, Gustavo Hernandez, Yuan Yu Lin, Adrian Marchetti

## Abstract

Wind-driven upwelling followed by relaxation results in cycles of cold nutrient-rich water fueling intense phytoplankton blooms followed by nutrient-depletion, bloom decline, and sinking of cells. Surviving cells at depth can then be vertically transported back to the surface with upwelled waters to seed another bloom. As a result of these cycles, phytoplankton communities in upwelling regions are transported through a wide range of light and nutrient conditions. Diatoms appear to be well-suited for these cycles, but their responses to them remain understudied. To investigate the bases for diatoms’ ecological success in upwelling environments, we employed laboratory simulations of a complete upwelling cycle with a common diatom, *Chaetoceros decipiens*, and coccolithophore, *Emiliania huxleyi*. We show that while both organisms exhibited physiological and transcriptomic plasticity, the diatom displayed a distinct response enabling it to rapidly shift-up growth rates and nitrate assimilation when returned to light and available nutrients following dark, nutrient-deplete conditions. As observed in natural diatom communities, *C. decipiens* highly expresses before upwelling, or frontloads, key transcriptional and nitrate assimilation genes coordinating its rapid response to upwelling conditions. Low iron simulations showed that *C. decipiens* is capable of maintaining this response when iron is limiting to growth, whereas *E. huxleyi* is not. Differential expression between iron treatments further revealed specific genes used by each organism under low iron availability. Overall, these results highlight the responses of two dominant phytoplankton groups to upwelling cycles, providing insight into the mechanisms fueling diatom blooms during upwelling events.

## Introduction

Phytoplankton communities in coastal upwelling regions disproportionately contribute to global primary production and serve as the base of highly productive food webs (1). Intermittent upwelling of cold, nutrient-rich water from depth fuels this productivity with blooms that eventually deplete essential inorganic nutrients and sink to depth as upwelling subsides. As a result of upwelling and relaxation, phytoplankton undergo a cycle in which subsurface populations are vertically transported with upwelled water masses to seed surface blooms and return to depth as upwelling subsides and nutrients are depleted (2–4). Herein this loop is referred to as the upwelling conveyer belt cycle (UCBC; Fig. 1A).

**Fig. 1.**
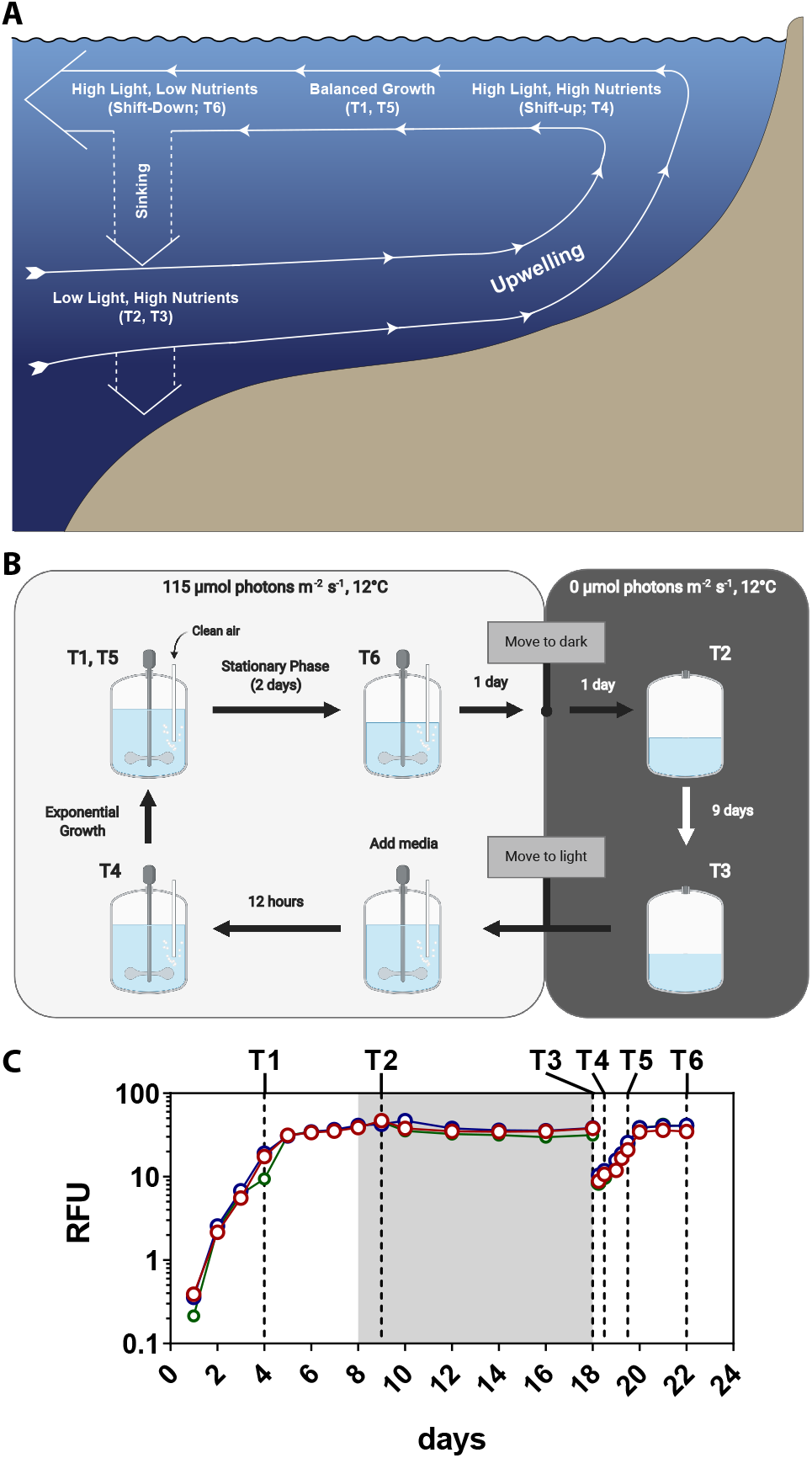
(A) Conceptual overview of the upwelling conveyer belt cycle (UCBC) reproduced from Wilkerson and Dugdale (5) and annotated with experimental time points that most closely represent different UCBC stages. Importantly, nutrients were not supplied at T2 and T3 resulting in a low light, low nutrient scenario. (B) Schematic of the experimental design to closely mimic different UCBC stages. (C) Growth curves in raw fluorescence units (RFUs) for C. decipiens in the high iron treatment throughout the UCBC simulation. The 10-day dark period is denoted with a grey background. Sampling time points are annotated with dotted lines. Data for the remaining experiments are shown in Fig. S1.

The physiological response of phytoplankton to the up-welling portion of the UCBC is referred to as the shift-up response and includes an acceleration of processes such as nitrate uptake, assimilation, and growth (2, 5). Diatoms are believed to be particularly suited for shift-up leading to their dominance of phytoplankton blooms following up-welling (4, 6, 7). When upwelling delivers cells and nutrients into well-lit surface waters, diatoms quickly respond to available nitrate and increase their nitrate uptake rates compared to other phytoplankton groups allowing them to bloom (8). This phenomenon may partially be explained by frontloading nitrate assimilation genes, or maintaining high gene expression leading up to an upwelling event such that nitrate assimilation transcripts are already abundant once upwelling conditions return (7). In addition, diatoms uniquely integrate their nitrogen and carbon metabolic pathways enabling a rapid response (9).

Coupled to rapid nitrate uptake rates, phytoplankton in natural communities have been observed to dramatically alter their carbon-to-nitrogen (C:N) ratios (7, 8, 10). C:N ratios well above the predicted Redfield value (6.6:1) imply that cells faced nitrogen limitation as the upwelled waters aged and then sank to depth, but this alteration has yet to be shown in the context of a complete UCBC cycle and without the influence of other non-phytoplankton detrital material that would affect these measurements (7, 8, 10). Ultimately, such shifts in cellular elemental quotas associated with sinking cells can decouple the biogeochemical cycling of these elements.

The responses of phytoplankton within the UCBC are also influenced by the availability of the micronutrient iron. In the California and Peru/Humboldt Upwelling Zones for example, iron delivery is primarily dependent on riverine input and upwelling-driven resuspension of continental shelf sediments (11–15). In areas with steep continental shelves, reduced interaction between upwelled waters and sediment results in decreased iron relative to macronutrients and iron limitation as the phytoplankton bloom develops. As up-welling is anticipated to intensify from climate change, increased upwelled nitrate has the potential to be further unmatched by upwelled iron causing an expansion of iron limitation regions (1, 16). Furthermore, ocean acidification may cause reduced iron bioavailability to phytoplankton (17, 18).

An understanding of phytoplankton responses to the UCBC remains limited and is primarily based on a relatively small collection of field studies focused on the physiological shift-up response resulting in an incomplete view of the cycle (4, 8). Importantly, potential differences in responses among phytoplankton groups may influence both the community structure and biogeochemical cycling in upwelling areas. Iron limitation may alter phytoplankton responses to the UCBC but also are uncharacterized (1).

Using a combined physiological and transcriptomic approach, we examine the responses of phytoplankton throughout the various light and nutrient conditions associated with the UCBC via culture-based simulations (Figs. 1B and 1C). Cultures were grown in exponential phase until nutrient exhaustian and stationary growth. After three days in stationary phase, cultures were moved to darkness for 10 days to simulated sinking out of the euphotic zone. Following this dark period, a portion of the culture was transferred to fresh medium and returned to light to simulate upwelling until stationary growth was again observed. The simulations included subsampling that was performed immediately before and 12 hours after the upwelling portion of the UCBC to evaluate the ability of cells to exhibit a shift-up response. Low iron simulations were included to evaluate responses under iron-limitation.

In order to compare two phytoplankton lineages commonly present in upwelled waters, a representative diatom, *Chaetoceros decipiens*, and coccolithophore, *Emiliania huxleyi*, were used. Both isolates were recently obtained from the California Upwelling Zone. The genus *Chaetoceros* is considered the most abundant and diverse diatom genus and is highly prevalent in the California coastal waters (19, 20). *E. huxleyi* is a globally distributed dominant bloom-forming coccolithophore and found to be one of the most abundant coccolithophores within a coastal California time series (21–23). Together, this comparison highlights changes in phytoplankton C:N ratios which influence biogeochemical cycling of these elements and examines unique physiological and molecular responses that provide insight into phytoplankton blooms in upwelling regions.

## Results and Discussion

### Physiological and transcriptomic responses through-out the UCBC

*C. decipiens* and *E. huxleyi* displayed clear physiological responses to the different UCBC conditions under iron-replete conditions (Fig. 2; black bars). Both reduced chlorophyll *a* under stationary growth phases (T2 and T3; Figs. 2A, 2B, and S1). Photosynthetic efficiency (F_v_:F_m_) was also lower at the stationary phase time points (T2 and T3) compared to the initial exponential growth time point (T1)(Figs. 2C and 2D). In *C. decipiens*, these declines were significant (P < 0.001, Dataset S1). Following the dark period, both species increased photosynthetic efficiency when the cells returned to exponential growth which is consistent with previous field observations (7).

**Fig. 2.**
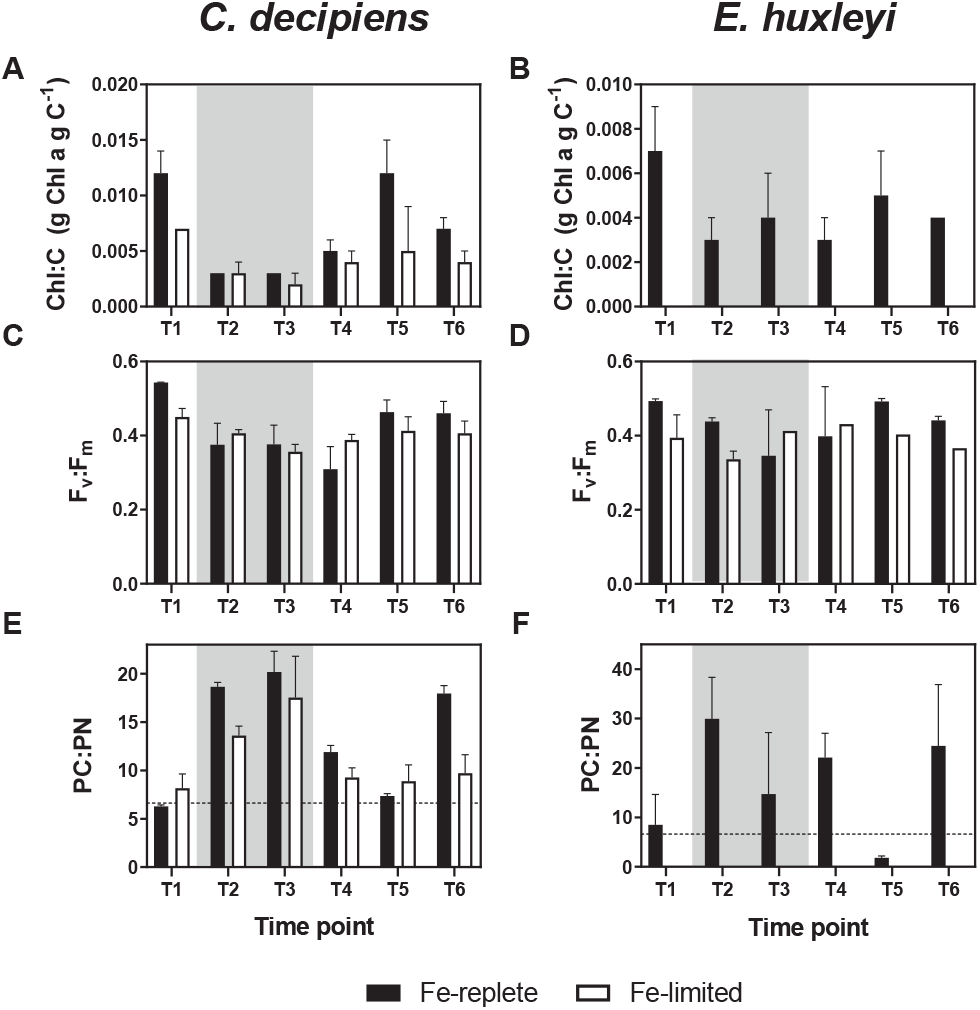
Biochemical and photophysiological measurements for *C. decipiens* and *E. huxleyi* in the iron-replete (black) and iron-limited (white) UCBC simulations: (A, B) Chlorophyll *a* to carbon ratios (g Chl *a* g C^−1^). (C, D) Maximum photochemical yields of photosystem II (F_v_:F_m_). (E, F) Particulate carbon to particulate nitrogen ratios. The dashed line indicates the Redfield C:N ratio (106:16). Grey background shading indicates measurements for time points in the dark. Error bars indicate the standard deviation of the mean (n = 3). Data are not available in *E. huxleyi* for chlorophyll *a* and for PC:PN at all time points in the iron-limited cultures.

Cellular C:N ratios for both species also fluctuated throughout the cycle (Figs. 2E and 2F). Both species were growing near Redfield-predicted values (6.6:1) during the initial exponential growth phase and increased their C:N ratios during stationary phase following N depletion (Fig. S2). As with chlorophyll *a* and (F_v_:F_m_) in *C. decipiens*, these differences were highly significant (P < 0.0001, Dataset S1). Previous field observations of particulate C:N from phytoplankton communities at depth have also shown values far exceeding the Redfield ratio reaching as high as 60:1, although uncertainty remained as to whether these results were a result of cellular modifications or simply C-rich detrital material (7, 8, 10). With a maximum C:N of 20.2 ± 2.1 for *C. decipiens* at T3 and 29.96 ± 8.4 for *E. huxleyi* at T2 under the iron-replete conditions, it can be inferred that a large proportion of the high carbon relative to nitrogen can be attributed to cells suffering from N and/or light limitation although in *E. huxleyi* these data also include inorganic carbon that is incorporated into coccoliths (Fig. S2). Similarly under N-limitation, several *Thalassiosira* species were shown to have C:N ratios over 20 which can be attributed to an accumulation of carbohydrates (24). Previous laboratory experiments that mimicked sinking cells with the diatom *Pseudo-nitzschia australis* grown in excess N relative to Si did not show as dramatic of an increase in C:N suggesting that Si limitation, as can often occur in conjunction with iron-limited diatoms in the California coastal upwelling region, may influence these elemental stoichiometries (25, 26).

To examine gene expression profiles in both organisms throughout the UCBC, transcriptome-wide co-expression was analyzed with Weighted Gene Co-Expression Network Analysis (WGCNA). WGCNA resulted in 23 modules (clusters of interconnected genes) for *C. decipiens* and 58 modules for *E. huxleyi*. Significant positive correlations (Pearson; P < 0.05) between modules and all time points apart from T6 for *C. decipiens* were found indicating that there is distinct and coordinated expression of certain genes within each UCBC stage for both organisms (Figs. 3 and S3). Several modules were significantly associated with the time point pairs that represented similar environmental conditions (T1 and T5; T2 and T3). Eighteen modules in *C. decipiens* and forty modules in *E. huxleyi* had significant positive correlations with each time point. Importantly, these significant associates were largely different between time points indicating that both organisms exhibited a high degree of transcriptional responsiveness to the different UCBC conditions.

**Fig. 3.**
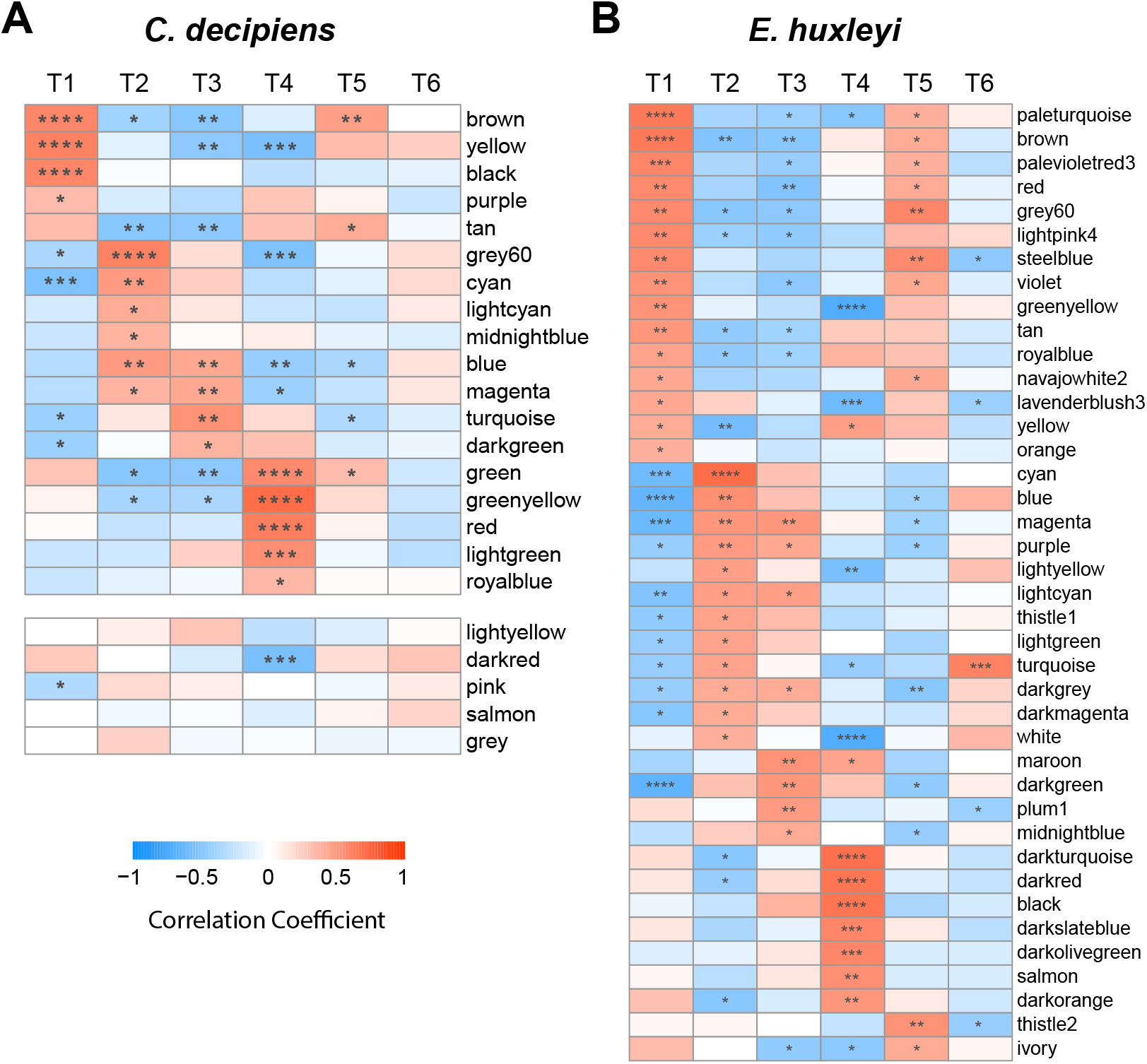
WGCNA module-time point correlations (Pearson) for (A) *C. decipiens* and (B) *E. huxleyi* at the time points throughout the UCBC. Significance in correlation is shown within each cell as follows: *P ^*≤*^ 0.05, **P ^*≤*^ 0.01, ***P ^*≤*^0.001, ****P ^*≤*^ 0.0001. Row names indicate WGCNA modules. Modules with no postitive significant correlations are shown separated in panel A for *C. decipiens* and in Fig. S3 for *E. huxleyi*.

Genes associated with each UCBC stage were identified by first examining genes within modules that correlated with each time point and then by determining which genes within those modules were also correlated with each time point (Fig. 3, Datasets S2 and S3). By comparing KEGG orthologs (KOs) and protein family (Pfam) annotations of contigs associated at each time point, similarities or differences in expression for both organisms throughout the UCBC were examined. Overall, *C. decipiens* and *E. huxleyi* showed highly divergent responses throughout the UCBC. When solely comparing KOs and Pfams detected in both transcriptomes, 513 unique KOs and 518 unique Pfams were significantly associated at any time point in both species; however, most of these were not associated with the same time points in both organisms. Furthermore, more KOs or Pfams were significantly associated with one of the organisms rather than both at each time point (Table 1).

**Table 1.** The number of KEGG Orthologs and protein families (Pfam) that were significantly associated with *C. decipiens, E. huxleyi*, or both (shared) at each time point. To be considered, the KEGG ortholog and Pfam must have been in both reference transcriptomes. Significance was determined as being both correlated individually with a time point and being in a WGCNA module correlated with a time point (Pearson). Significance cutoffs of Benjamini & Hochberg adjusted P-values < 0.05 were used.

|  |  | T1 | T2 | T3 | T4 | T5 | T6 |
| --- | --- | --- | --- | --- | --- | --- | --- |
| KEGG | Shared | 66 | 22 | 4 | 109 | 14 | N/A |
|  | *C. decipiens* | 166 | 98 | 320 | 411 | 38 | N/A |
|  | *E. huxleyi* | 300 | 133 | 39 | 295 | 142 | 52 |
| Pfam | Shared | 100 | 44 | 23 | 142 | 16 | N/A |
|  | *C. decipiens* | 145 | 83 | 250 | 320 | 39 | N/A |
|  | *E. huxleyi* | 312 | 190 | 58 | 263 | 146 | 103 |

Broadly, this divergence in responses represents differing metabolic investments and potential niche partitioning between diatoms and coccolithophores; however, some similarities can be observed by examining specific KEGG orthologs, KEGG modules, and Pfams associated with each organism at each time point (Datasets S2 and S3). In both organisms, many photosynthesis related genes were associated with time points when light was available. Expression of glycolysis and fatty acid biosynthesis genes during exponential growth was also common. At 12 hours after returning to the light, both highly expressed genes associated with translation and amino acid biosynthesis as they were in transition to resume growth.

In examining KOs found in both organisms that were correlated with different time points, differences in the timing of transcription-related gene expression were observed (Fig. 4, Datasets S2 and S3). These include RNA polymerases and spliceosome genes to generate mature RNA, and TRAMP and exosome complexes that can interact to degrade abnormal rRNAs and spliced-out introns (27). *C. decipiens* displayed elevated expression of these gene sets during stationary growth, particularly at T3, compared to its exponential growth phases. In an opposite pattern, *E. huxleyi* decreased expression of these genes during stationary phase and elevated expression during exponential growth and at T4 in response to a return to light and nutrients. WGCNA indicated that these gene sets were significantly associated with T3 in *C. decipiens* but with T4 in *E. huxleyi* (Datasets S2 and S3).

**Fig. 4.**
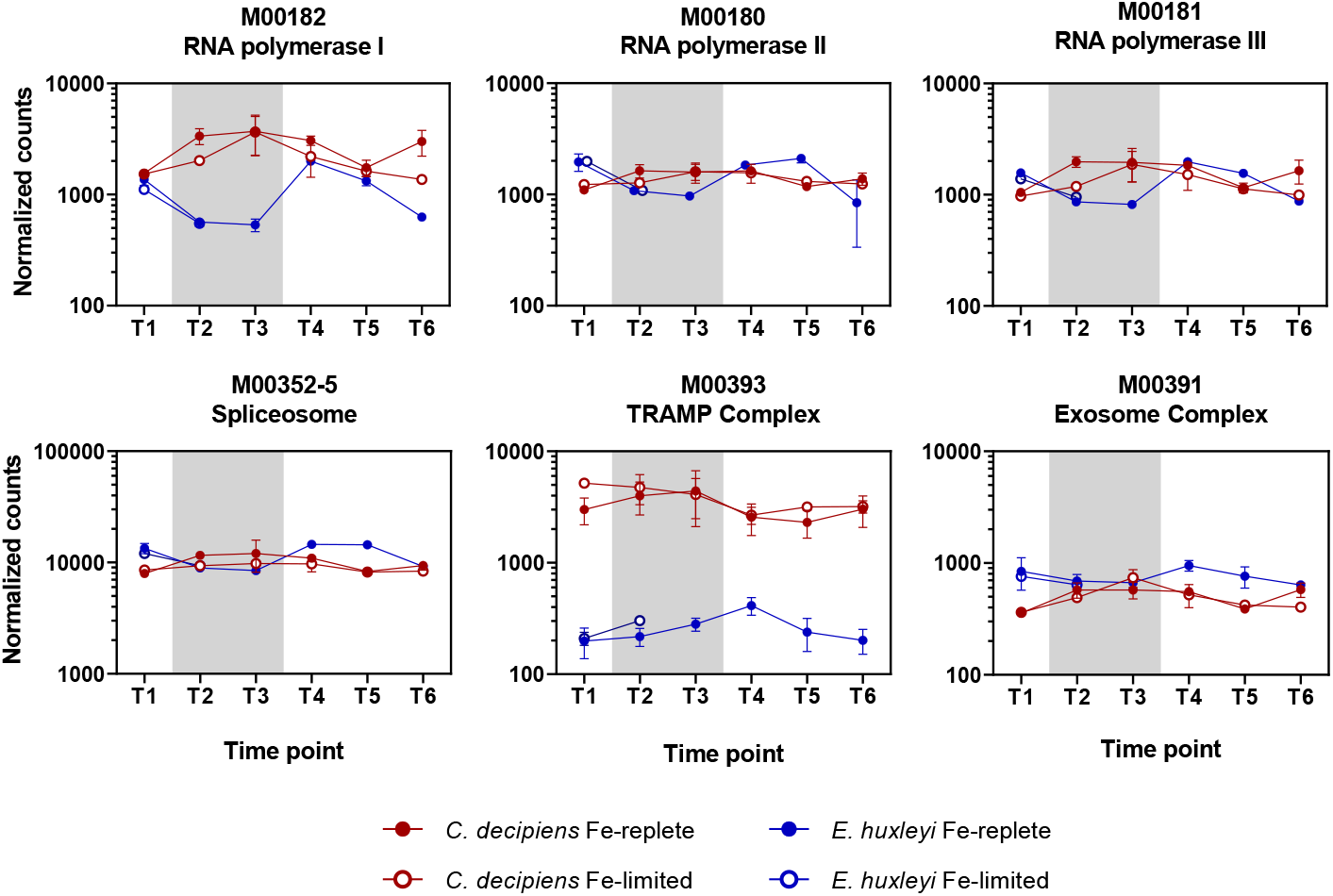
Summed DESeq2-normalized counts assigned to select transcription-related KEGG modules (M) in *C. decipiens* (red) and *E. huxleyi* (blue) under Fe-replete (closed circles) and Fe-limited (open circles) treatments. Error bars indicate the standard deviation of the mean (n= 3).

These opposing patterns indicate relatively high cellular investments in transcription during differing growth phases. In *C. decipiens*, this higher expression corresponds to periods when conditions are not optimal for growth, i.e., a lack of light and nutrients, whereas higher expression corresponds to periods leading to or within exponential growth in *E. huxleyi*. This difference in timing of expression may drive transcriptional differences observed in the field where diatoms were found to highly express, or frontload, nitrogen assimilation genes prior to upwelling conditions in contrast to other groups including haptophytes (7). An increase in transcriptional machinery may be linked to the frontloading response by enabling an accumulation of nitrogen assimilation transcripts that ultimately assists in diatoms to outcompete other groups by being transcriptionally proactive rather than reactive in response to the relative rapid and frequent changes in environmental conditions (28).

### The shift-up response and nitrogen assimilation

Frontloading of nitrogen assimilation genes and an acceleration of nitrate uptake, or the shift-up response, is hypothesized to be linked to diatom success during the upwelling portion of the UCBC (2, 5, 8). As a result, the upwelling portion of the UCBC simulation was sampled after 12 hours (T4) to investigate the physiological capability for relatively rapid shift-up. By comparing T3 through T5, a view from pre-upwelling, 12 hours after upwelling, and a return to exponential growth can be examined (Fig. 1).

A rapid shift-up response was observed in *C. decipiens* compared to *E. huxleyi. C. decipiens* returned to exponential growth within 24 hours of returning to light and nutrients whereas *E. huxleyi* did not respond appreciably for approximately 48 hours (Fig. S4). UCBC simulations with varying dark phases suggest that the capability to respond in *E. huxleyi* is impacted by the length of the dark phase with an inability to reinstate growth after 20 days in the dark (Fig. S5). Concurrently, the C:N ratio in *C. decipiens* rapidly declined from 20.18 ± 2.13 to 11.89 ± 0.70 within the first 12 hours of returning to light and nutrients suggesting that *C. decipiens* rapidly assimilated nitrate once returned to optimal growth conditions (Figs. 2E and 2F). After 36 hours in the light, cellular C:N ratio in *C. decipiens* returned to the Redfield ratio. In contrast, *E. huxleyi* maintained a high C:N ratio (22.10 ± 4.91) after returning to light for 12 hours, but the C:N ratio dramatically declined to 1.80 ± 0.40 at 3 days after returning to the light when the cells reached exponential growth (Figs. 2F and S1). This initial lack of decline indicates that *E. huxleyi* has a slower response to the newly available nitrate compared to *C. decipiens*; however, once the cells reach exponential growth, *E. huxleyi* may exhibit luxury uptake and store excess nitrate prior to an increase in cell division as evidenced by the later decline in their C:N ratio in conjunction with the highest cellular nitrogen quotas (Fig. S6). Altogether, these results align with the relatively rapid shift-up response of diatoms compared to other phytoplankton groups, and that nitrate uptake rates are likely rapidly increasing to result in the observed changes.

Results from a field-based simulated upwelling experiment suggest that frontloading of nitrogen assimilation genes is an underlying molecular mechanism behind these responses (7). Diatoms were observed to highly express nitrate assimilation genes [nitrate transporter (*NRT*), nitrate reductase (*NR*), nitrite reductases (*NiR*)] prior to upwelling events in order to be transcriptionally proactive, or frontload, to rapidly assimilate nitrate and outcompete other phytoplankton groups once conditions are optimal for growth.

In comparing the relative and summed expression of nitrogen metabolism genes in *C. decipiens* and *E. huxleyi* from the UCBC simulation, *C. decipiens* exhibited this distinct and proactive frontloading response whereas *E. huxleyi* did not (Figs. 5A and 5B). *C. decipiens* significantly upregulated the aforementioned nitrogen assimilation genes during stationary phase (P < 0.05) with highest expression at 10 days in the dark, i.e., the simulated pre-upwelling condition. WGCNA also placed these genes in the blue module which was significantly associated with the dark stationary phase time points (Fig. 3, Dataset S2). Upon returning to high light and nutrients, expression of these genes significantly declined and returned to previous levels observed during exponential growth (P < 0.01, Fig. 5B). Diatoms also possess two nitrite reductases (Nir) both of which are localized to the chloroplast; however, one utilizes ferredoxin (Fd) and the other utilizes NAD(P)H (9). Under exponential growth, *Fd-Nir* transcript abundance was higher than *NAD(P)H-Nir*; however, under stationary growth in the light and dark, *NAD(P)H-Nir* transcript abundance was higher suggesting potential metabolic advantages of each nitrite reductase under these different growth conditions (Fig. 5A).

**Fig. 5.**
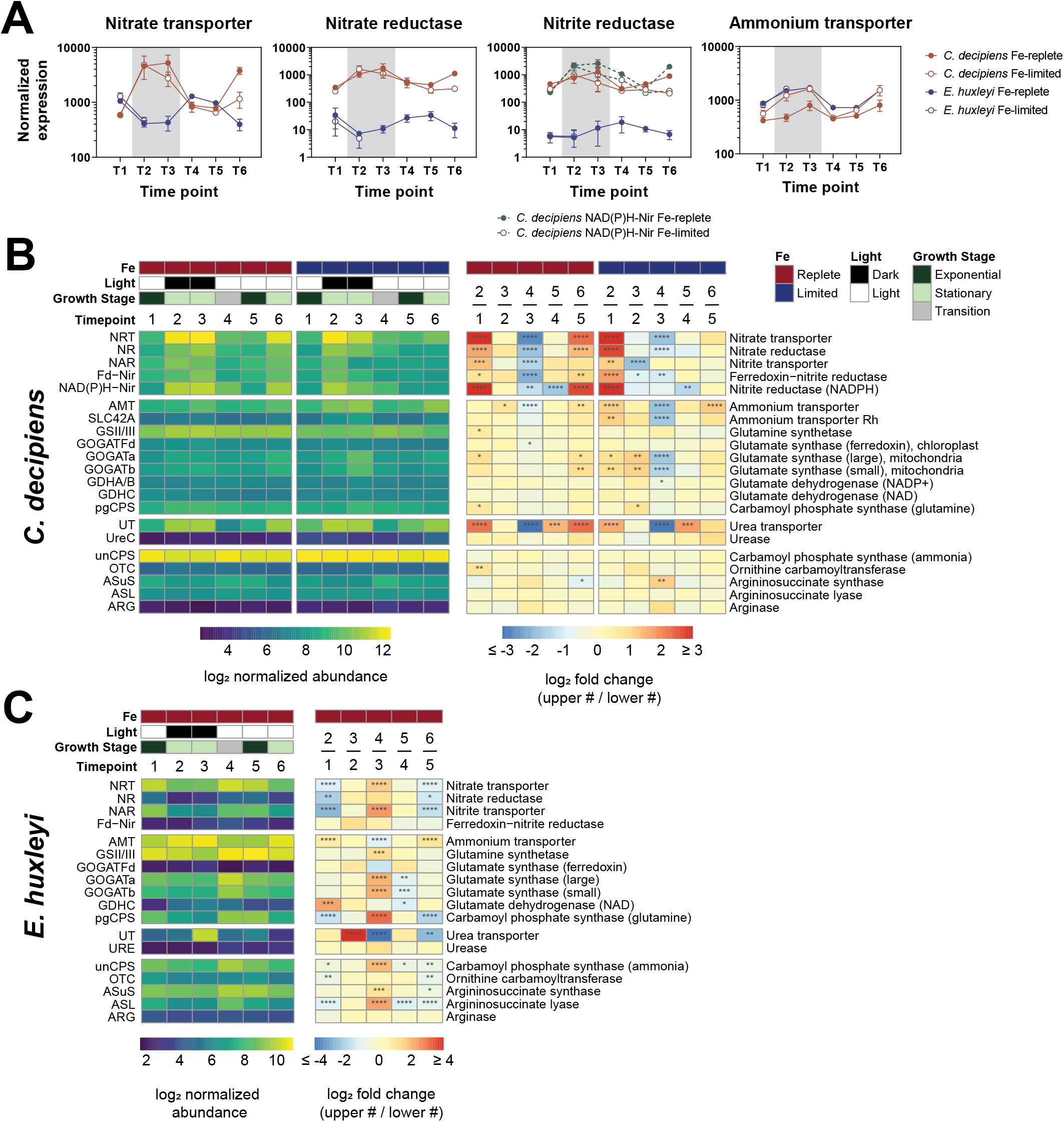
Expression of nitrogen assimilation and utilization genes throughout the UCBC. (A) Line graph view of averaged nitrate assimilation and ammonium transporter normalized counts in both *C. decipiens* (red) and *E. huxleyi* (blue) under Fe-replete (closed circles) and Fe-limited conditions (open circles). *C. decipiens* possesses two nitrite reductases; therefore, Fd-Nir is shown with a solid red line and NAD(P)H-Nir is shown with a dashed green line. Error bars indicate the standard deviation of the mean (n = 3). (B) Heatmaps showing both the log_2_ normalized abundance and log_2_ fold change values throughout the UCBC. Iron status, light level, growth stage, and time point are annotated above the heatmaps where appropriate. Fold changes are compared by examining one time point to the next (later time point / earlier time point) and annotated as the upper numbered time point / the lower numbered time point. Significance in differential expression between time points is shown within each cell as follows: *P ^*≤*^ 0.05; **P ^*≤*^ 0.01; ***P ^*≤*^ 0.001; ****P ^*≤*^ 0.0001. (C) Corresponding heatmaps for *E. huxleyi* as described for panel A. Iron-limited experiments are omitted due to a lack of replication in RNA for all time points.

In contrast, *E. huxleyi* displayed an opposing expression pattern for their nitrate assimilation genes (Fig. 5A). During stationary growth, *E. huxleyi* significantly downregulated transcripts for nitrate transporters and nitrate reductase (P < 0.01); nitrite reductase expression remained similar across time points (Fig. 5B). Once returned to the light and nutrients (T4), *E. huxleyi* significantly upregulated transcripts for nitrate transporters (P < 0.0001) in response to nitrate availability and optimal growth conditions. Nitrate reductase and nitrite reductase expression also remained low compared to other *E. huxleyi* transcripts (16^th^-40^th^ percentile) throughout the experiment; therefore, in conjunction with low cellular C:N ratios, these data support that *E. huxleyi* may store nitrate for some time post-upwelling.

Unlike the nitrate assimilation genes, ammonium transporters (*AMT*) and urea transporters (*UT*) displayed a similar transcriptional pattern in both organisms; however, while nitrate transporters were among the most highly expressed nitrogen metabolism genes in *C. decipiens* (87^th^-100^th^ percentile), *E. huxleyi* highly expressed ammonium transporters (94^th^-100^th^ percentile) despite ammonium not being added to the seawater medium. For *AMT*, these patterns were driven by the mostly highly expressed copies in both organisms with some copies displaying different gene expression patterns (Fig. S7). High expression of *AMT* and *UT* during stationary phase suggests that both organisms are tuned to exploit released ammonium and urea as nitrate was depleted, although urease expression remained low in both (4^th^-21^st^ percentile; Figs. 5B, 5C, and S7). *E. huxleyi* has also previously been shown to prefer ammonium over nitrate (29). The high total level of expression of *AMT* relative to the expression of other genes in *E. huxleyi* and compared to that of *AMT* expression in *C. decipiens* corroborates this preference for ammonium and implies that the two species exhibit different nitrogen preferences.

The divergent responses between the two species with frontloading observed in the diatom is increasingly apparent when examining the nitrogen metabolism genes downstream of the assimilation genes, the GS-GOGAT [glutamine synthetase (*GS*) and glutamate synthase (*GOGAT*)] cycle and the ornithine-urea cycle (OUC; Figs. 5B, 5C and S8). The GS-GOGAT cycle catalyzes ammonium into organic nitrogen. The ornithine-urea cycle (OUC) has been shown to aid the model diatom *Phaeodactylum tricornutum* in returning from N-limiting conditions by serving as a hub for nitrogen redistribution and pathway to satisfy arginine demand (9, 30). *E. huxleyi* also possesses an OUC which is hypothesized to serve a similar role (31).

In *C. decipiens*, significant differences in expression between time points were nearly absent in both the GS-GOGAT cycle under the iron-replete conditions and OUC genes under both iron treatments (Fig. 5B). These results correspond to previous field observations of constitutive and relatively high expression in diatoms pre- and post-upwelling such that the cell already possesses the cellular machinery to channel assimilated nitrogen towards growth (7). Like the nitrate assimilation genes, *E. huxleyi* exhibits a different transcriptional response for the GS-GOGAT and OUC genes in response to the changing UCBC conditions. OUC genes were generally downregulated under stationary growth and upregulated in response to returning to light at T4 (P < 0.05). In response to returning to light, *E. huxleyi* strongly upregulated GS-GOGAT genes (P < 0.001; Fig. 5C). Both *C. decipiens* and *E. huxleyi* displayed low and constitutive expression of arginase and urease (3^rd^-31^st^ percentile), the final steps of the OUC pathway which similarly were not shown to exhibit a frontloading response in the same degree in the field (Fig. S9)(7). The OUC thus likely serves to satisfy demand for arginine in both organisms as predicted through metabolic flux models in *P. tricornutum* (9). Much like the nitrate assimilation genes, *C. decipiens* appears to frontload the GS-GOGAT and OUC genes whereas *E. huxleyi* exhibits a more responsive transcriptional pattern.

N-limitation has previously been studied extensively in several diatoms and *E. huxleyi* providing points of comparison. In the model pennate diatom *P. tricornutum*, genes encoding nitrate transporters, nitrate reductases, and nitrite reductases were upregulated under N limitation (32, 33), although a study examining these genes in a short-term response associated them with nitrogen replete conditions (9). Upregulation of these genes under N starvation in most cases was further shown in *Fragilariopsis cylindrus, Pseudonitzschia multiseries*, and several *Skeletonema* species (33, 34); therefore, most studies are consistent with our findings here of upregulation under N limitation suggesting that this response is shared among diatoms.

For GS-GOGAT and OUC-related genes, a pattern in gene expression is less apparent. Most studies reported changes in expression of these genes under N-replete or - deplete conditions in contrast to the lack of change observed here in *C. decipiens*, although the directionality of the expression among studies is not consistent (9, 32, 33). However, in *F. cylindrus, P. tricornutum*, and *P. multiseries*, stable expression of OUC genes was shown between N conditions apart from the first step, carbamoyl-phosphate synthase, similar to observations in this study (33). Therefore, unlike nitrate assimilation genes, transcriptional regulation of these genes appears to be variable among diatoms as previously suggested (33). The coordinated frontloading response of GS-GOGAT and OUC genes with the nitrate assimilation genes may be a strategy more exclusively employed by bloom-forming diatoms residing in upwelling environments.

Aligning with the results presented here, *E. huxleyi* has previously been shown to downregulate or show reduced activity of NRT, NR, and NiR while upregulating *AMT* under N-limitation, although these results are also not entirely consistent across studies (29, 31, 35–37). *E. huxleyi* also was shown to downregulate the initial OUC genes under N limitation, the one exception here being arginosuccinate synthase at T2 (31). GS-GOGAT gene expression was similarly lower in previous studies under N limitation albeit not significantly different here (31).

### Influences of iron limitation on UCBC responses

Iron bioavailability is an important control on phytoplankton growth in upwelling areas characterized by narrow continental shelves (1). Biogeochemical modeling of the California Current Ecosystem indicates that iron limitation is highly prevalent although only occurring at small scales and short durations relative to N limitation (38). This iron limitation occurs in both the surface mixed layer and at depth as shown with subsurface chlorophyll maximum layers (13, 39). Furthermore, projected increases in upwelled nitrate unmatched by iron coupled to ocean acidification is projected to expand iron limitation in these regions (1, 17, 18). Here UCBC simulations were performed under iron-limiting conditions to examine the effects of iron status on responses throughout the cycle.

Within the first exponential phase, the growth rate in *C. decipiens* declined from 0.86 ± 0.05 d^−1^ in iron-replete cells to 0.36 ± 0.08 d^−1^ in iron-limited cells (Fig. S1). The reduction in growth rates was much smaller for *E. huxleyi* (0.68 ± 0.05 d^−1^ to 0.63 ± 0.08 d^−1^). Reductions in F_v_:F_m_ were also observed in both organisms indicating iron stress in the low iron treatments. With F_v_:F_m_ already impacted by iron stress, the variation as a result of light and/or nitrogen stress was comparatively less to when iron was replete. Previous experiments have also suggested that *E. huxleyi* is well-suited to grow under low Fe concentrations (40). Higher growth rates may suggest that *E. huxleyi* has a competitive advantage under Fe-limitation, yet during the UCBC, iron-limited *E. huxleyi* cells were unable to reinstate exponential growth upon returning to light in two out of three replicates, indicating that *E. huxleyi* can be severely impacted by both the combination of prolonged darkness and Fe-limitation (Fig. S4). Importantly, in the one *E. huxleyi* bottle that was able to grow, it did not return to exponential growth for 48 hours while *C. decipiens* appeared to immediately grow similar to the observations in the iron-replete cells. This observation is in line with diatoms being able to outcompete coccolithophores under dynamic conditions (41).

Iron limitation can influence nitrate assimilation as the required enzymes, nitrate and nitrite reductase, can account for greater than 15 percent of the cellular iron requirement (42, 43). Iron limitation has also been shown to influence the expression of the genes that encode these proteins (44, 45). In *C. decipiens*, C:N ratios followed a similar pattern when comparing the iron-replete to iron-stressed conditions (Fig. 2). In early stationary phase (T2 and T6), cellular C:N ratios were significantly reduced relative to iron-replete cells at the same time points, but reached similar values after 10 days in the dark. Importantly, C:N ratios in iron-limited cells still quickly approached the Redfield ratio upon returning to the light (T4) indicating that the shift-up response in Nassimilation may not be significantly impacted by low iron conditions in *C. decipiens*.

Iron limitation altered the expression levels of certain N-related genes at specific time points in *C. decipiens* (Fig. S10). The nitrate assimilation pathway was downregulated at T3 and T6. This decreased expression at T3 made expression levels between T3 and T4 similar which may explain similar responses observed pre- and post-upwelling in the field-based incubations where iron stress in the initial community may have occurred (7).

Genes encoding ammonium and urea transporters as well as urease were upregulated under low iron as expected with nitrate assimilation being impacted. Under low iron, variability in expression between time points increased for mitochondrial GOGAT genes compared to the fairly stable expression in the iron-replete treatment with significant up-regulation under low iron at T3 (Figs. 5B and S10). The mitochondrial GOGAT genes are specialized to recover recycled nitrogen and/or assimilate urea (9); therefore, in conjunction with the ammonium and urea transporters being up-regulated under low iron, this transcriptional response is consistent with a greater emphasis on utilizing recycled forms of N. As with the lack of change in OUC genes throughout the UCBC, there were no significant differences in OUC transcript abundances between iron treatments at all time points.

Other differentially expressed genes show how *C. decipiens* responds to iron limitation during the UCBC (Fig. 6A, Dataset S4). Similar to other diatoms, several transcriptional indicators of iron stress were significantly upregulated in the iron-limited treatment. Transcripts for the inorganic iron uptake gene phytotransferrin (*pTF*; previously known as *ISIP2a*) were upregulated at all time points in iron-limited cells (P < 0.0001)(17). Flavodoxin (FLDA) is an iron-free replacement for the photosynthetic electron transfer protein ferredoxin (46), and diatoms can possess multiple copies of *FLDA* with some that are not iron responsive (47). Two copies of *FLDA* were detected, one of which was significantly upregulated at all time points in iron-limited cells (P < 0.0001) and the other was not. A gene encoding for a metacaspase similar to one identified in T. pseudonana (*TpMC4*), was significantly upregulated at all time points (P < 0.0001); this protein has been implicated in programmed cell death suggesting a triggering of this process in iron-limited cells (48).

**Fig. 6.**
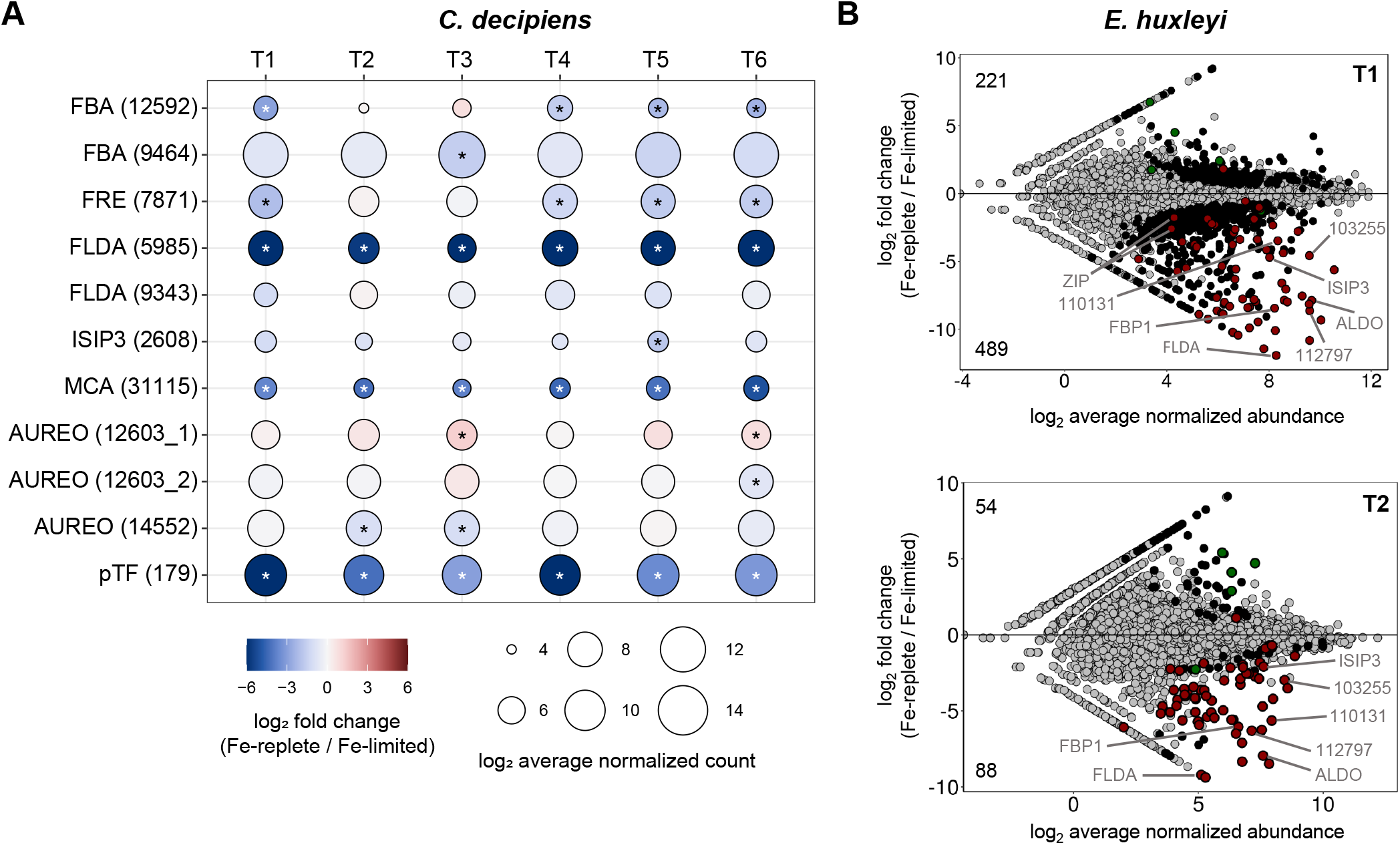
Iron-related gene expression (A) Selected genes in *C. decipiens*. Genes with contig IDs in parentheses are shown on the vertical axis. Time points are shown on the horizontal axis. Circle size and color correspond to the log_2_ average DESEq2-normalized abundance and the log_2_ fold change as shown in the legend. Contigs that were significantly upregulated under iron-limiting conditions are denoted with an asterisk (P < 0.05). Genes are abbreviated as follows: FBA fructose-bisphosphate aldolase, FRE ferric reductase, FLDA flavodoxin, ISIP3 iron-starvation induced protein 3, MCA metacaspase, AUREO aureochrome, pTF phytotransferrin. (B) MA plots showing differentially expressed contigs in E. huxleyi between iron treatments at T1 (top) and T2 (bottom). Contigs are colored to denote differences in significant expression (P < 0.05): gray is not significant at the displayed time point, black is significant at the displayed time point but not in the other, green is significant in both with a positive fold change in the other time point, and red is significant in both with a negative fold change in the other time point. The total number of significantly differentially expressed genes with positive or negative fold changes are shown in the top left or bottom left respectively. Genes are abbreviated as follows: FBA fructose-bisphosphate aldolase, FBP1 ferrichrome-binding protein, FLDA flavodoxin, ISIP3 iron-starvation induced protein 3, ZIP ZRT, IRT-like protein. Numbers used for gene annotation correspond to protein IDs in the *E. huxleyi* CCMP1516 genome.

Some genes displayed differential expression only at certain time points throughout the UCBC. *ISIP3*, a gene of unknown function normally highly expressed under iron limitation in diatoms, was only upregulated at T1 (P = 0.06) and T5 (P = 0.005) indicating that this gene may not be a reliable marker for iron stress in *Chaetoceros* under serial limitation or colimitation scenarios unlike *T. oceanica* (49–51). A ferric reductase in the cytochrome b_5_ family was differentially expressed only in the light (P < 0.001). This contig is homologous to *FRE3* and *FRE4* in *P. tricornutum* which are speculated to be involved in heme-based iron uptake (44). Two class I fructose 1,6-bisphosphate aldolase (*FBA*) contigs were upregulated under low iron, although one was only up-regulated in the light and other in the dark (T3)(P < 0.05). These enzymes are important for cellular C metabolism and are commonly found to be upregulated in iron-limited diatoms (45, 52).

Three contigs encoding aureochromes were differentially expressed as a function of iron status at certain time points. Aureochromes are light-induced transcription factors specific to stramenopiles and all three of these in *C. decipiens* contain the characteristic bZIP domain for DNA binding and the light-oxygen-voltage (LOV) domain for blue-light sensing (53, 54). These proteins have also previously been implicated in diel regulation of gene expression (55, 56). None were differentially expressed between iron treatments during exponential growth. Two (12603_2 and 14552) increased in expression under iron-limitation when in stationary phase in the dark, suggesting that these genes have an increased role in regulating gene expression when both nitrogen and iron are limiting, a colimitation scenario which may be commonplace in the ocean (57). The third aureochrome (12603_1) was up-regulated in the high iron treatment under stationary growth suggesting it has a more pronounced role in gene regulation when iron is replete but nitrogen is limiting.

Despite its prevalence and ability to grow under low iron, differential expression in *E. huxleyi* between varied iron treatments has not been analyzed previous to this study. This analysis was restricted to T1 and T2 as only these time points had replicates available (Fig. 6B, Dataset S5). More genes were significantly differentially expressed at T1 compared to T2 (Fig. 6B). Only two contigs showed significant opposing regulation between T1 and T2, that is, one was upregulated at one time point but downregulated at the other.

Many of the genes upregulated under iron limitation were similar to those that have been frequently observed in diatoms. These include the previously mentioned genes *ISIP3, FBA Class I*, and *FLDA*. High expression of *FBP1* and ZIP family divalent metal transporters were also detected. *FBP1* is involved in siderophore-based Fe uptake in diatoms suggesting a similar functional role in *E. huxleyi* (58). ZIP family transporters may also permit transport of ferrous iron (59). Other highly expressed genes have no annotated function (e.g., *E. huxleyi* CCMP1516 IDs: 103255, 110131, 112797) suggesting that *E. huxleyi* possesses or employs unique mechanisms for coping with iron limitation. These genes and the proteins they encode are clear targets for future study and possible markers of iron stress for *E. huxleyi* in the natural environment.

### Conclusions

Phytoplankton in upwelling regions are exposed to highly dynamic conditions as a result of upwelling cycles. In these upwelling cycle simulations, both a representative diatom, *C. decipiens*, and the coccolithophore, *E. huxleyi*, displayed physiological and transcriptional plasticity to these cycles. Part of this physiological response included transitioning to high cellular C:N ratios which has previously been observed in natural populations and influences the biogeochemical cycling of these elements. Although both phytoplankton species were transcriptionally responsive, their gene expression patterns at each time point was highly divergent. In particular, *C. decipiens* exhibited a pattern of frontloading transcriptional and nitrate assimilation genes, which may explain certain diatoms’ ability to rapidly respond and outcompete other groups once upwelling events occur. As observed in the diatom *Cylindrotheca fusiformis*, nitrate reductase transcripts accumulate under nitrogen starvation to enable extremely rapid translation into protein upon addition of nitrate (60); therefore, the diatom frontloading response observed here may be restricted to high transcription and rely on post-transcriptional regulation as the synthesis of mature proteins is expensive.

As a highly diverse group, diatoms likely occupy a wide range of niches and growth strategies (20). The response characterized here may be relatively exclusive to bloom-forming diatoms in upwelling regions although transcriptional responses in nitrate assimilation genes appear fairly consistent as indicated from other diatoms in laboratory studies. Furthermore, it is likely even among bloom-forming diatoms that there are further divergences in the transcriptional responses as observed in a previous field simulation (7). Additional study of other phytoplankton species, particularly other bloom-forming diatoms such as *Pseudo-nitzschia* and *Thalassiosira*, will be necessary to more fully understand the spectrum of responses to UCBC conditions.

Both *C. decipiens* and *E. huxleyi* employed mechanisms in response to iron stress that are common to responses previously observed in diatoms; however, certain genes such as aureochromes appear to only be differentially expressed when nitrogen and/or light is simultaneously limiting, indicating a unique response under colimitation scenarios. As Si and Fe limitation are often linked in diatoms (25, 61), examination of Si and Fe colimitation responses throughout the UCBC would provide further environmentally-relevant insights.

In *E. huxleyi*, many of the genes that were differentially expressed under iron limitation are not functionally characterized. Importantly, *E. huxleyi* was unable to respond to prolonged darkness under iron-limiting conditions whereas *C. decipiens*’s shift-up response once returned to the light was unaffected by iron status. Some projections suggest increased iron limitation in upwelling environments as a result of ocean acidification and increases in upwelled nitrate relative to iron (1, 17); however, these findings suggest that diatoms will continue to exhibit the shift-up response and out-compete other phytoplankton groups under both high and low iron upwelling conditions.

Clearly, there are a number of differences between the simulation experiments presented here and the conditions found in the natural environment (Fig. 1). Temperature and pressure would not be constant as they were in the simulations. Upwelled waters are also more acidic, and pH adjustments were not made (62). Although nutrient concentrations would increase as the cells sink to depth, nitrate, the depleted nutrient, was not supplemented until the cells returned to light following the dark phase. In addition, iron in nature is bound to a wide variety of organic ligands and supply is dynamic whereas our simulations utilized steady-state buffered conditions with iron bound to a single chelator, EDTA (63). The laboratory cultures also contained significantly higher cell densities compared to those found in the natural environment, and the phytoplankton strains studied individually here, albeit non-axenically, prevented these responses from being examined with direct interactions and competition among a larger number of species. Such complexities will require further field-based observations and experimentation to more fully understand phytoplankton responses to the UCBC.

## Materials and Methods

### Experimental design

Two phytoplankton species isolated from the California Upwelling Zone were used in this study: a diatom, *Chaetoceros decipiens* (UNC1416), and a coccolithophore, *Emiliania huxleyi* (UNC1419). Species isolation and identification are described in Lampe et al. (7). Cultures were grown in artificial Aquil* medium following trace metal clean (TMC) techniques at 12°C and continuous light at approximately 115 µmol photons m^−2^ s^−1^ (64–66). Starting macronutrient concentrations were modified from standard Aquil* amounts such that nitrate rather than silicate would be drawn down and depleted first by the phytoplankton isolates (50 µmol L^−1^ NO_3_, 10 µmol L^−1^ PO_4_, 200 µmol L-1 H_4_SiO_4_). The culture media was chelexed in a TMC room, microwave sterilized, allowed to cool, and then supplemented with filter-sterilized (0.2 µm Acrodisc®) EDTA-trace metals (minus iron) and vitamins (B_12_, thiamine, and biotin). Trace metals were buffered using 100 µmol L^−1^ EDTA and added at the concentrations listed in Sunda, et al. (66) except Cd which was added at a total concentration of 121 nmol L^−1^. Premixed Fe-EDTA (1:1) was added separately at a total concentration of 1370 nmol L^−1^ for the high iron treatments or 3.1 nmol L^−1^ for the low iron treatments. The resulting concentrations of iron not complexed to EDTA (Fe’) were estimated as 2730 pmol L^−1^ (high iron) and 6 pmol L^−1^ (low iron)(66) resulting in iron-replete and iron-limited growth, respectively. Vitamins were added at f/2 concentrations. Media were allowed to equilibrate overnight before use.

Cultures were acclimated to media at the two iron concentrations using the semi-continuous batch culture technique in acid-cleaned 28 mL polycarbonate centrifuge tubes until growth rates were consistent between transfers. Cultures were then grown in a 1 L polycarbonate bottle, and when in exponential phase, 5 mL (high iron) or 20 mL (low iron) of each culture were then transferred to triplicate clean 2 L polycarbonate bottles fitted with Teflon tubing for sub-sampling and bubbling. Seawater was continuously stirred and bubbled with air passed through 1.2 mol L^−1^ HCl then through TMC Milli-Q before entering the culture bottles to remove any potential contaminants in the air. Cultures were not axenic although sterile techniques were used for all culture work to minimize contamination.

The upwelling conveyer belt cycle (UCBC) was simulated by growing or transitioning the cultures to varying light and nutrient conditions (Fig. 1B, Fig. S1). First, to simulate bloom and termination phases, cultures were grown in exponential phase until nutrient exhaustion, at which time cells entered stationary phase. After three days in stationary phase, bubbling and stirring were halted and the bottles were moved to a dark incubator (0 µmol photons m^−2^ s^−1^, 12°C) to simulate sinking out of the euphotic zone (2, 3, 7). Although temperature, pressure, and nutrient concentrations would change in deeper waters in the natural environment, temperature and pressure were kept constant and cells were not supplied with additional nutrients during this stage. After 10 days, 500 mL of culture was transferred into 1.5 L of fresh medium and returned to the light with stirring and bubbling resumed. Cultures were subsequently grown until stationary growth was measured for two days, thus completing the cycle. Subsampling was always performed at the same time of day (approximately 09:00 Eastern Time) except T4 which occurred 12 hours after T3 (21:00).

The 10-day time frame was selected based on our previous field-based simulated upwelling experiments where satellite data indicated that upwelling had not occurred in the region for approximately 13 days although intervals are certainly variable in nature (7). Additionally, preliminary iron-replete UCBC experiments suggested that *E. huxleyi* is incapable of reinstating growth after 20 days of darkness and thus unable to complete the UCBC simulation with that duration of darkness (Fig. S5). This provided an upper limit for the length of the dark period further leading to the selection of 10 days.

Subsampling was performed by first dispensing into sterilized bottles under a laminar flow hood in a TMC room, then immediately aliquoting from this subsample. Relative abundances and growth throughout the experiment were assessed by regular measurements of blank-corrected raw fluorescence units (RFUs) with a Turner 10-AU fluorometer. Specific growth rates were calculated during exponential growth phases from the natural log of RFUs versus time (67).

### UCBC dark phase duration experiments

Experimental trials were conducted with both phytoplankton isolates to determine the effect of different lengths of darkness on growth during UCBC simulations (Figs. S5). The two isolates were grown in acid-cleaned 28 mL polycarbonate centrifuge tubes with synthetic Aquil* medium that contained the same macronutrient and iron concentrations as the iron-replete treatment. The dark phase was varied with five-day intervals between five and twenty days in duration. After the dark phase was complete, 7 mL of culture was transferred into 21 mL of fresh media. Changes in phytoplankton biomass and calculation of growth rates were assessed with near-daily measurements of blank-corrected raw fluorescence units (RFUs) using a Turner 10-AU fluorometer (67).

### Cell counts

Five mL samples were preserved in 2% Lugol’s iodine in glass vials. Cells were then enumerated on an Olympus CKX41 inverted microscope using a 1 mL Sedgwick-Rafter counting chamber after allowing the cells to settle for five minutes and counting a minimum of 300 cells within 10-30 fields of view.

### Chlorophyll

Fifty mL of culture was filtered through a 0.45 µm mixed cellulose ester filter (25 mm; EMD Millipore Corporation, Burlington, MA, USA) under gentle vacuum pressure (< 100 mm Hg) and immediately frozen at −80°C until analysis. Chlorophyll a extraction was performed using 90% acetone at −20°C for 24 h and measured via in vitro fluorometry on a Turner Designs 10-AU fluorometer using the acidification method (68).

### Particulate carbon and nitrogen

Particulate carbon (PC) and nitrogen (PN) were obtained by gentle vacuum filtration of 50 mL of culture onto a pre-combusted (450°C for 6 hours) GF/F filter. Filters were immediately stored in petri dishes at −20°C. Prior to analysis, filters were dried at 65°C for 24 hours then encapsulated in tin. Total nitrogen and carbon were quantified with a Costech 4010 CHNOS Elemental Combustion system according to U.S. Environmental Protection Agency Method 440.0 (69). Three blanks were run alongside the samples and were all below the detection limits (0.005 mg N and 0.071 mg C). Samples were not acidified and include particulate inorganic carbon including that from the coccolith in *E. huxleyi*. Carbon and nitrogen on a per cell basis were calculated by dividing the particulate carbon or nitrogen concentration by cell concentration.

### F_v_:F_m_

The maximum photochemical yield of Photosystem II (F_v_:F_m_) was measured using a Satlantic FIRe (Fluorescence Induction and Relaxation System) (70, 71). Samples were acclimated to low light for 20 min prior to measuring the minimum (F_o_) and maximum (F_m_) fluorescence yields. Data were blank corrected using microwave-sterilized Aquil medium. The resulting F_v_:F_m_ was derived from the induction profile using a saturating pulse (Single Turnover Flash; 20,000 µmol photons m^2^ s^−1^) for a duration of 100 µs. The gain was optimized for each sample (400, 600, or 800), and the average of 50 iterations was obtained.

### Dissolved nitrate and nitrite concentrations

Filtrate from the 0.45 µm filters used for RNA was transferred to polypropylene tubes and immediately frozen at −80°C. Dissolved nitrate + nitrite (NO_3_ + NO_2_) concentrations were quantified with an OI Analytical Flow Solutions IV auto analyzer according to EPA method 353.4 (72). The detection limit for NO_3_ + NO_2_ was 0.2 µmol L^−1^.

### RNA Extraction

Approximately 300 mL of culture was filtered onto 0.45 µm Pall Supor® polyethersulfone filters (47 mm) using gentle vacuum pressure and immediately frozen and stored at −80°C. For the *Chaetoceros* experiments, total RNA was extracted using the RNAqueous-4PCR Total RNA Isolation Kit (Ambion, Foster City, CA, USA) according to the manufacturer’s protocol with an initial bead beating step to disrupt cells. For the *E. huxleyi* experiments, total RNA was extracted using TRIzol reagent (Invitrogen, Carlsbad, CA, USA) according to the manufacturer’s protocol except for an initial bead beating step and two instead of one chloroform steps to separate proteins and DNA. Trace DNA contamination was removed by DNase 1 (Ambion) digestion at 37°C for 45 min, and RNA was purified with the RNeasy MinElute Cleanup Kit (Qiagen, Germantown, MD, USA). Quality and quantity were checked on 1% agarose gels and with a Nanodrop 2000 spectrophotometer (Thermo Fisher Scientific, Waltham, MA, USA).

### Reference transcriptomes

Phytoplankton cultures were grown to late exponential phase and extracted as described under RNA Extraction. RNA libraries were created with either the Illumina TruSeq Stranded mRNA Library Preparation Kit for *C. decipiens* or the KAPA Stranded mRNA-Seq kit for Illumina platforms for *E. huxleyi*. Samples were sequenced on an Illumina MiSeq (300 bp, paired-end reads) and an Illumina HiSeq 2500 with one lane in high output mode (100 bp, paired-end reads) and another lane in rapid run mode (150 bp, paired-end reads)(Table S1).

Raw reads were trimmed for quality with Trimmomatic v0.36 (73) then assembled *de novo* with Trinity v2.5.1 with the default parameters for paired-reads and a minimum contig length of 90 bp (74). Contigs were clustered based on 99% similarity using CD-HIT-EST v4.7 (75) and then protein sequences were predicted with GeneMark S-T (76). Protein sequences were annotated by best-homology (lowest E-value) with the KEGG (Release 86.0)(77), UniProt (Release 2018_03)(78), and PhyloDB (v1.076)(79) databases via BLASTP v2.7.1 (E-value ≤ 10-5) and with Pfam 31.0 (80) via HMMER v3.1b2 (Table S2). KEGG Ortholog (KO) annotations were assigned from the top hit with a KO annotation from the top 10 hits (https://github.com/ctberthiaume/keggannot).

### RNA library preparation and sequencing

RNA libraries for the UCBC simulations were constructed using a custom protocol for 3’ poly-A-directed mRNA-seq (also known as TagSeq) based on Meyer et al. (81) and adapted for Illumina HiSeq based on Lohman et al. (82) and Strader et al. (83). For most samples 1 µg but as low as 250 ng of total RNA were fragmented by incubating at 95°C for 15 min.

First strand cDNA was synthesized with SMARTScribe Reverse Transcriptase (Takara Bio, Mountain View, CA, USA), an oligo-dT primer, and template switching to attach known sequences to each end of the poly-A mRNA fragments. cDNA was then amplified with a PCR reaction consisting of 32 µL sterile H_2_O, 5 µL dNTPs (2.5 mM each), 5 µL 10X Titanium Taq Buffer (Takara Bio), 1 µL of each primer (10 µM) designed amplify the sequences attached during cDNA synthesis, and 1 µL of Titanium Taq DNA Polymerase (Takara Bio). The PCR was run with an initial denaturing step at 95°C for 5 min, then 17 cycles consisting of 95°C for 1 min, 63°C for 2 min, and 72°C for 2 min. cDNA amplification was then verified on a 1% agarose gel and purified with the QiaQuick PCR Purification Kit (Qiagen). cDNA was quantified with the Quant-iT dsDNA High-Sensitivity Assay (Invitrogen) and cDNA concentrations were then normalized to the same volume.

Samples were then barcoded with a PCR reaction consisting of 11 µL sterile H_2_O, 3 µL dNTPs (2.5 mM each), 3 µL 10X Titanium Taq Buffer (Takara Bio), 0.6 µL TruSeq Universal Primer (10 µM), 6 µL barcoding primer (1 µM), Titanium Taq Polymerase (Takara Bio), and 6 µL of purified cDNA with an initial denaturing step at 95°C for 5 min, then 5 cycles consisting of 95°C for 40 s, 63°C for 2 min, and 72°C for 1 min. Samples were barcoded from both ends using unique combinations of 6 bp sequences on both the TruSeq universal primer and barcoding primers. Products were then again confirmed on a 2% agarose gel and combined into small pools of 6-8 samples. The 400-500 bp region of these pools was extracted with the QIAquick Gel Extraction Kit (Qiagen), quantified with the Quant-iT dsDNA High-Sensitive Assay, and then mixed in equal proportions. The library was sequenced at the University of Texas at Austin Genomic Sequencing and Analysis Facility on Illumina HiSeq 2500 (three lanes, single-end 50bp reads) with a 15% PhiX spike-in to target approximately 8 million reads per sample (Table S2)(81).

### Gene Expression Analysis

PCR duplicates were removed based on a matching degenerate leader sequence incorporated during cDNA synthesis and a match of the first 20 bases as described by Dixon et al. (84). Reads were trimmed for adapter removal and quality with a minimum length of 20 bases with Trimmomatic v0.38 (73). Transcripts were quantified with Salmon v0.9.1 with the GC bias correction option against predicted proteins from *de novo* transcriptome assemblies generated via paired-end Illumina sequencing of the same strains (85). Normalized counts, fold change values, and associated Benjamini & Hochberg adjusted p-values were calculated with DESeq2 v1.22.2 (86). Normalized counts are based on the median of the ratios of the observed counts (87). Reference transcriptome and experimental results are further detailed in Tables S1 and S2 and Dataset S6.

Correlation network analysis of gene expression was performed using WGCNA using variance-stabilizing transformed counts generated by DESeq2 as input (88). All available samples from both the iron-replete and iron-limited treatments were used for this analysis (Dataset S6). Contigs with fewer than 10 normalized counts in more than 90% of the samples were removed before analysis as suggested by the WGCNA manual. A signed adjacency matrix was constructed for each organism using a soft-thresholding power of 14 for the *C. decipiens* samples and 22 for the *E. huxleyi* samples (Fig. S11). The soft-thresholding power was selected as the lowest power that satisfies the approximate scale-free topology criteria as described by Langfelder and Horvath (88). The adjacency matrix was then transformed into a Topological Overlap Matrix (TOM). Subnetworks, or modules, were created by hierarchical clustering of the TOM with the following parameters: minModuleSize = 30, deep-Split = 2. The grey module is reserved for genes that are not assigned to a module. The first principal component of each module, i.e. eigengene, was correlated (Pearson) with each time point and associated p-values were calculated. Contigs in modules that were significant for a specific time point were then also correlated (Pearson) to the same time points to evaluate significance on an individual contig basis (Dataset S2 and Dataset S3). The associated p-values were corrected for multiple testing using the Benjamini & Hochberg false discovery rate controlling procedure (89); significant correlations were those with an adjusted p-value less than 0.05. Parameters for all software used are listed in Table S3.

### Statistical Analyses

One- and two-way ANOVAs followed by Tukey’s multiple comparison test were performed on the biological and chemical properties of the seawater media in Graphpad PRISM v8.4.3.

### Data deposition

Raw reads for the *C. decipiens* and *E. huxleyi* experiments are deposited in the National Center for Biotechnology Information (NCBI) Sequence Read Archive (SRA) under the accession nos. SRP176464 and SRP178167 (Bioproject accession nos. PRJNA513340 and PRJNA514092) respectively. Raw reads for the reference transcriptomes are also deposited in SRA under the accession nos. SRP234548 and SRP234650 (Bioproject accession nos. PRJNA593314 and PRJNA593538). Assemblies and annotations for each reference transcriptome are deposited in Zenodo (doi:10.5281/zenodo.3747717 and doi:10.5281/zenodo.3747989). Example code for the DESeq2 and WGCNA analyses are available at https://github.com/rhlampe/simulated_upwelling_exps.

## ACKNOWLEDGEMENTS

We thank Liah McPherson, Jackie Millay, and Sharla Sugierski for assistance with the culture experiments and sample processing. This work was funded by National Science Foundation grants DGE-1650112, DGE-1650116 (Graduate Research Fellowship to R.H.L), OCE-1751805 (to A.M.), and ICER-1600506 (stipend for G.H.). Partial funding for sequencing was also provided by a Phycological Society of America Grant-in-Aid of Research (to. R.H.L.).

