## Supplementary Information for "Representative diatom and coccolithophore species exhibit divergent responses throughout simulated upwelling cycles"

**Robert H. Lampe[a*], Gustavo Hernandez[a], Yuan Yu Lin[a], and Adrian Marchetti[a#]**

[a]Department of Marine Sciences, University of North Carolina at Chapel Hill, Chapel Hill, NC, USA

**[#]For correspondence:**
Dr. Adrian Marchetti


*Present Addresses: Integrative Oceanography Division, Scripps Institution of Oceanography, University of California, San Diego, La Jolla, CA, USA; Microbial and Environmental Genomics, J. Craig Venter Institute, La Jolla, CA 92037, USA

**This file includes:**

Supplementary Figures 1-11
Supplementary Tables 1-3
Supplementary Data Set Information
References for SI Citations

**Fig. S1.** Cell counts and raw fluorescence units (RFU) of *C. decipiens* and *E. huxleyi* throughout the UCBC experiments under Fe-replete and Fe-limited conditions. The 10-day dark period is denoted with a grey background. Specific growth rates ($d^{-1}$) for both exponential growth phases are denoted as $\mu_1$ and $\mu_2$ respectively.

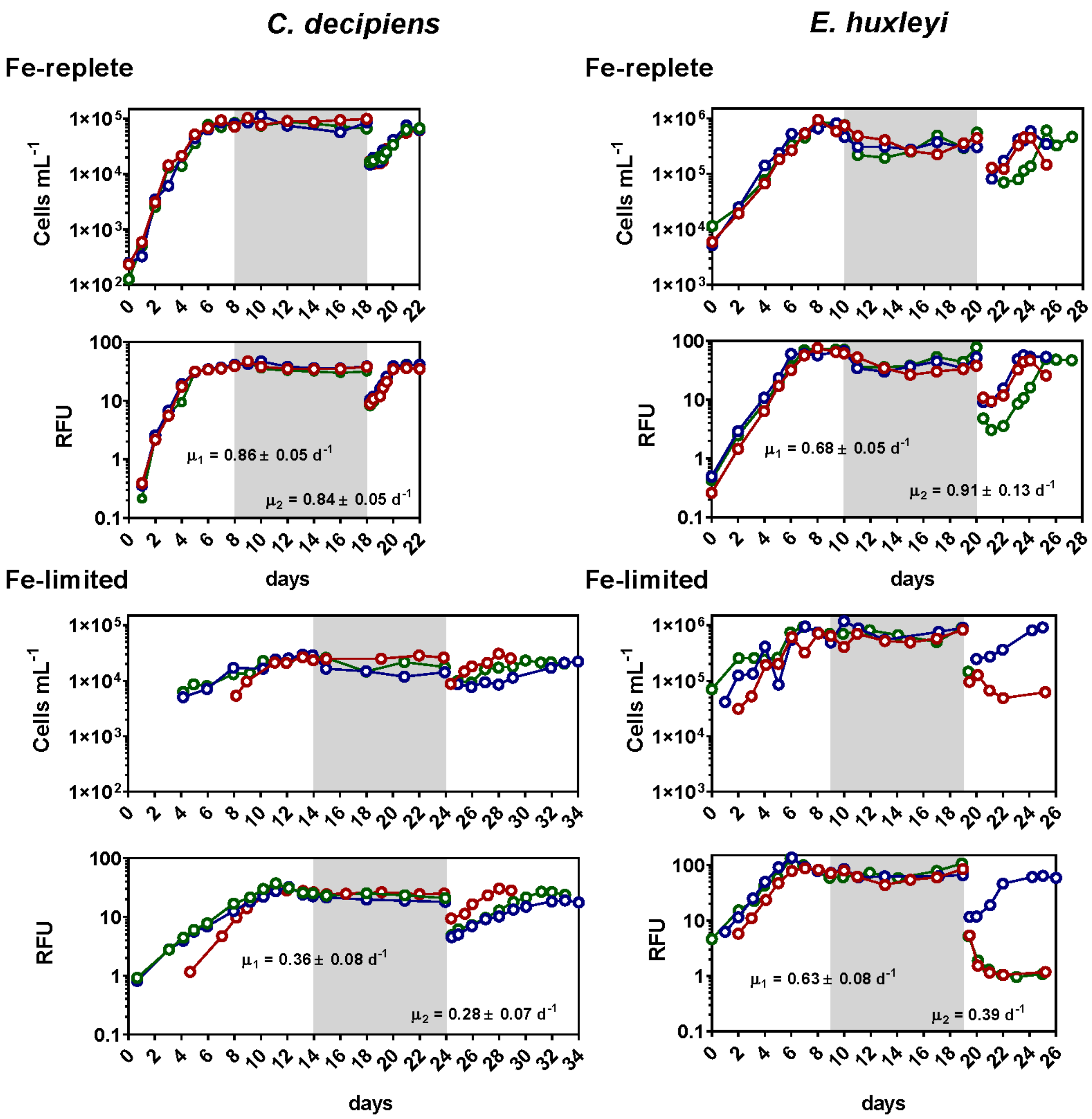

**Fig. S2.** Nitrate plus nitrite ($NO_3$+$NO_2$) concentrations throughout the UCBC simulations. Concentrations were not measured at T3 as no nitrate was added after T2. Only T1 and T2 were quantified for the iron-limited experiments in *E. huxleyi*. Gray background shading indicates measurements for time points in the dark.

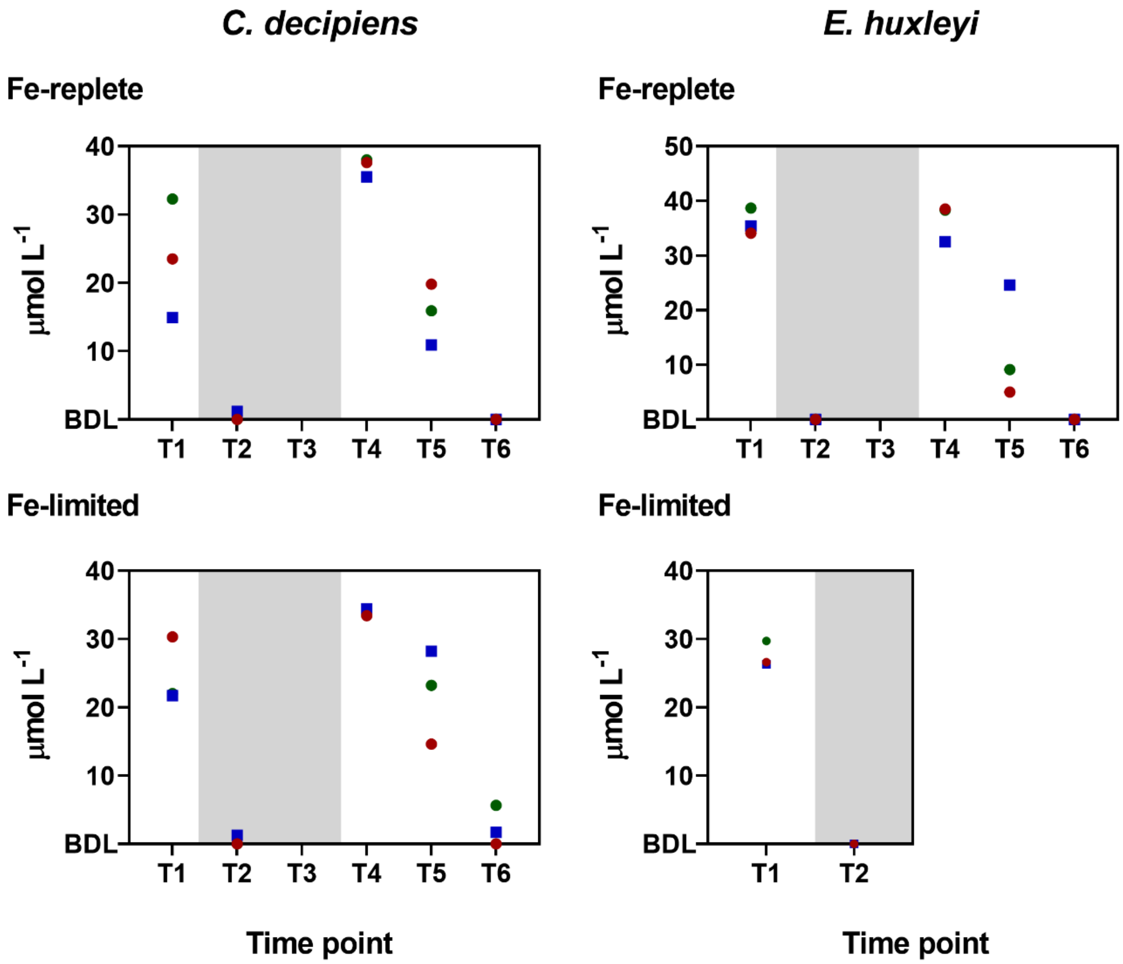

**Fig. S3.** WGCNA modules with no positive significant correlations with time points for *E. huxleyi*. Significance in correlation is shown within each cell as follows: *P ≤ 0.05, **P ≤ 0.01, ***P ≤ 0.001, ****P ≤ 0.0001.

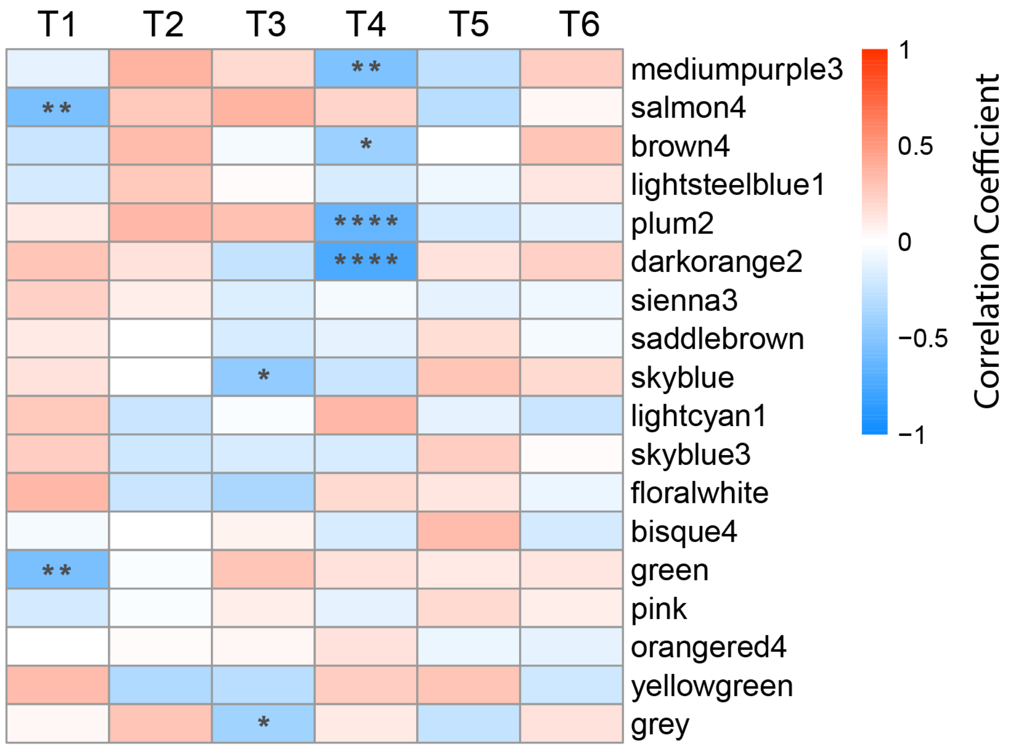

**Fig. S4.** Raw fluorescence units (RFU) of *C. decipiens* and *E. huxleyi* following the dark period under Fe-replete and Fe-limited conditions as shown in Fig. S1.

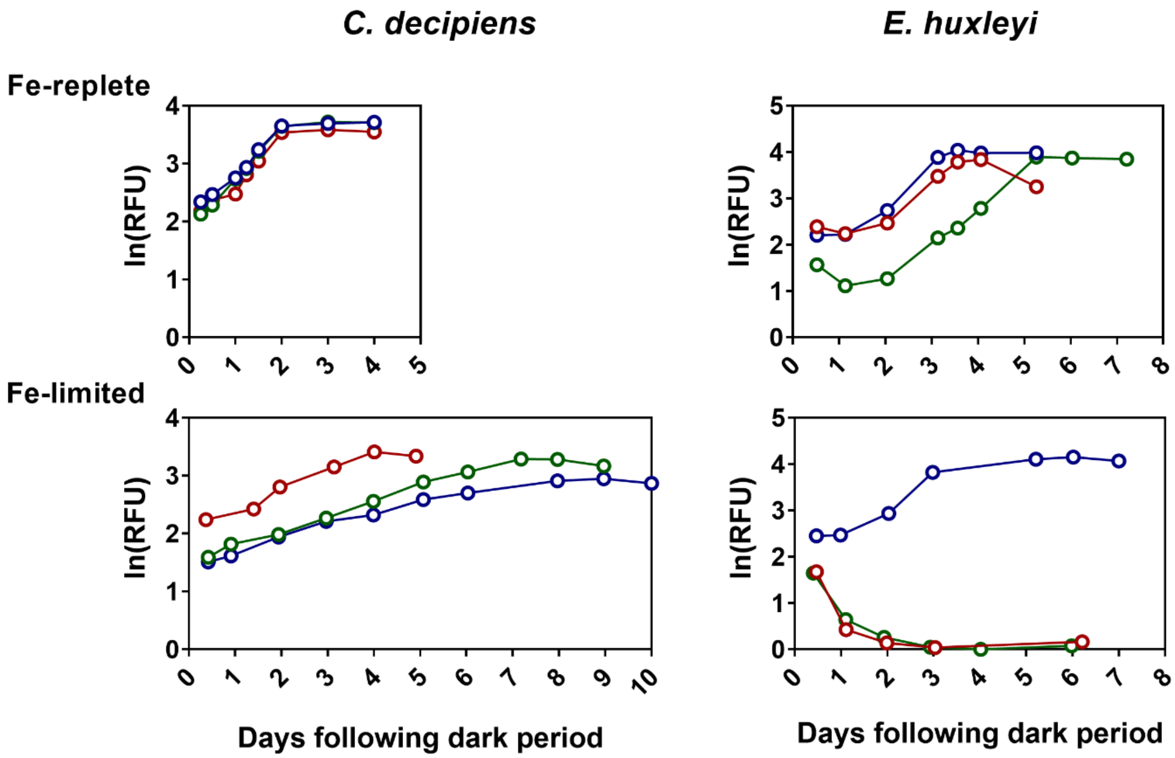

**Fig. S5.** Raw fluorescence units (RFUs) on a log scale from Fe-replete preliminary UCBC experiments (*Supplementary Materials and Methods).* Different dark periods were used with 5-day intervals and are denoted with a grey background.

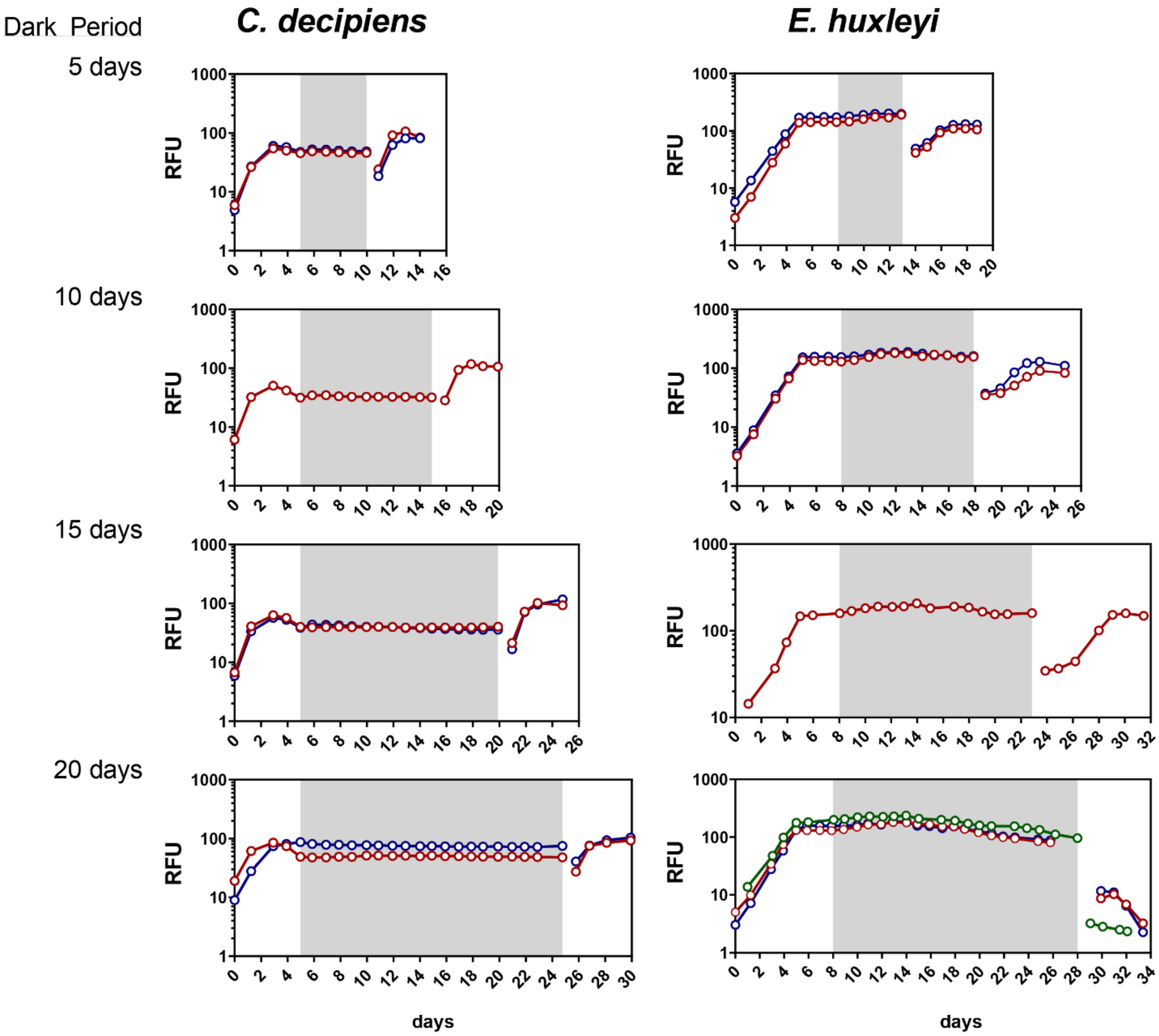

**Fig. S6.** N and C quotas on a per cell basis (pmol cell$^{-1}$) for (A, B) *C. decipiens* and (C, D) *E. huxleyi* in the Fe-replete (black) and Fe-limited (white) experiments. Data are not available in the low iron experiments for *E. huxleyi*. Gray background shading indicates measurements for time points in the dark. Error bars indicate the standard deviation of the mean (n = 3).

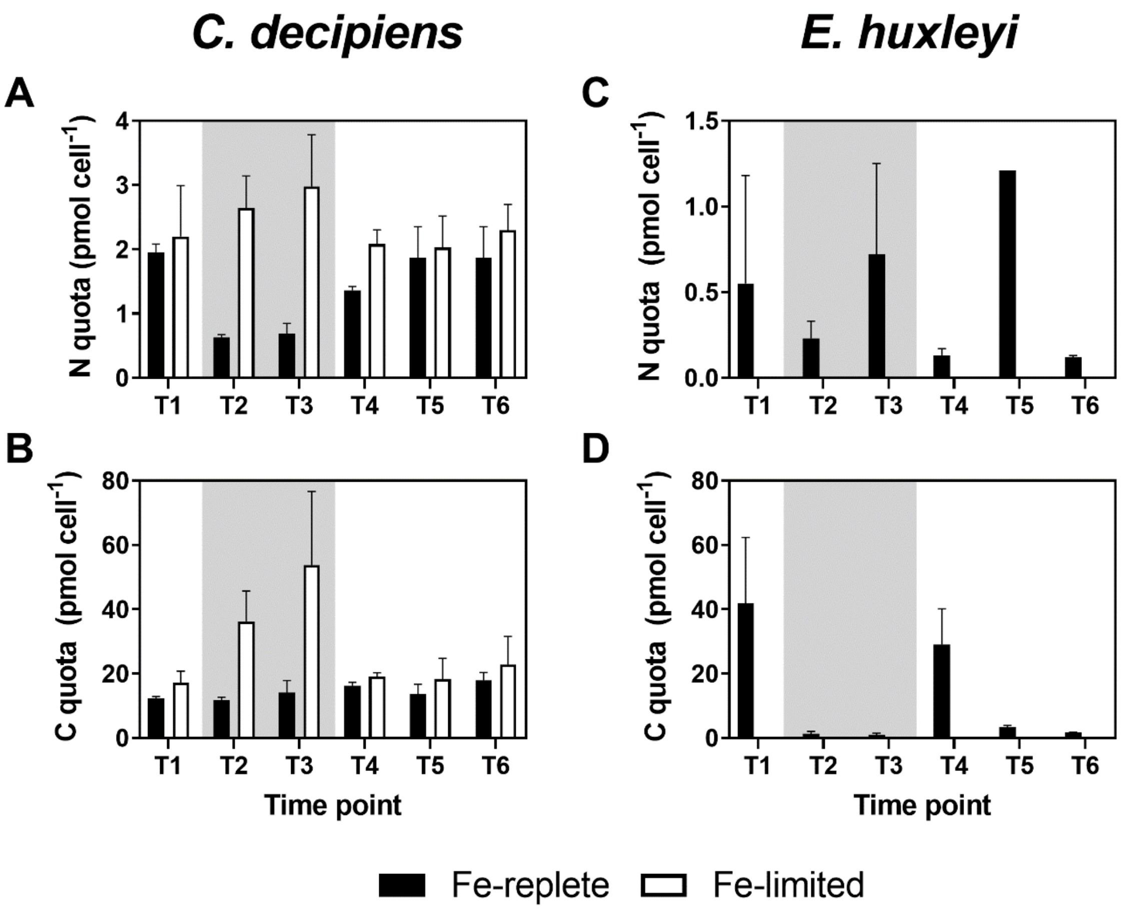

**Fig. S7.** Line graph view of DESeq2-normalized counts for individual *AMT* copies in *C. decipiens* (left) and *E. huxleyi* (right) under iron-replete (closed circles) and Fe-limited (open circles). Contig IDs are shown within each panel. Only contigs with an average normalized count greater than 50 across all samples are shown. Error bars indicate the standard deviation of the mean where n = 3 except *C. decipiens* iron-replete T5 and *E. huxleyi* iron-replete T3 where n = 2.

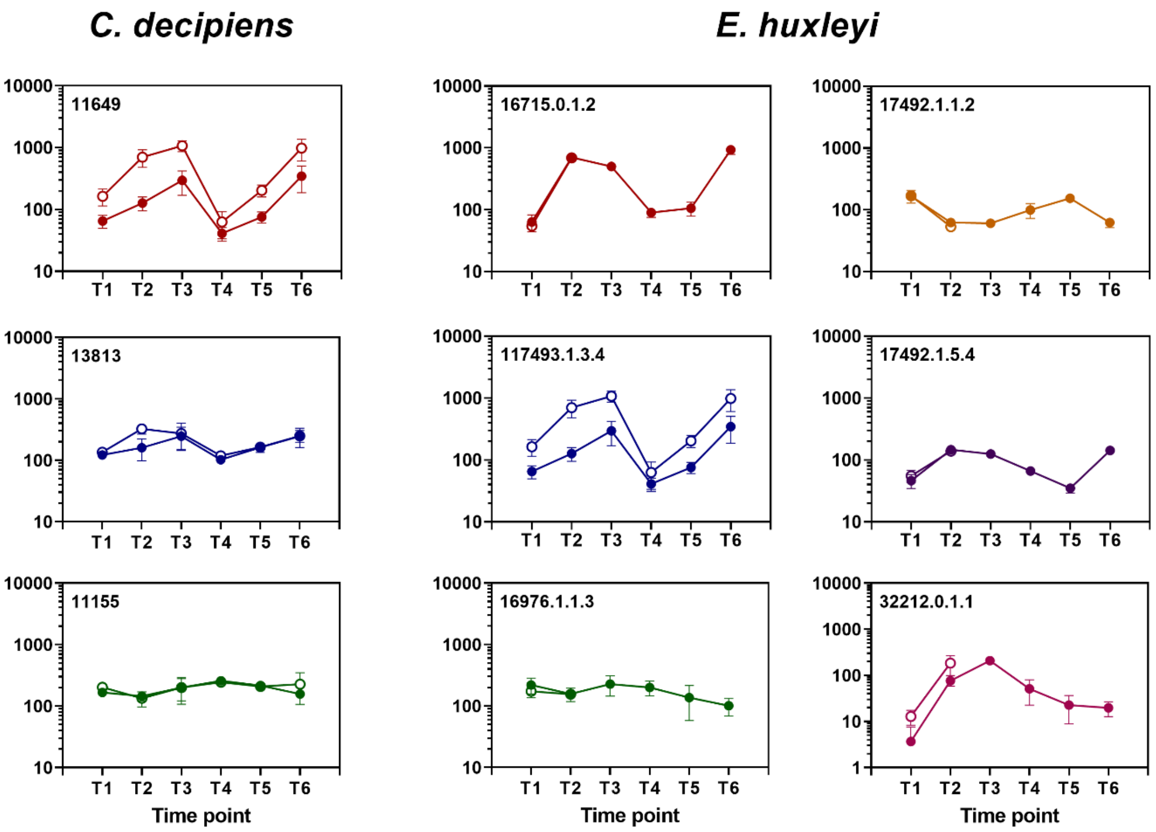

**Fig. S8.** Line graph view of GS-GOGAT DESeq2-normalized gene counts in both *C. decipiens* (red) and *E. huxleyi* (blue) under Fe-replete (closed circles) and Fe-limited (open circles) conditions. Gray background shading indicates measurements for time points in the dark. Error bars indicate the standard deviation of the mean where n = 3 except *C. decipiens* Fe-replete T5 and *E. huxleyi* iron-replete T3 where n = 2.

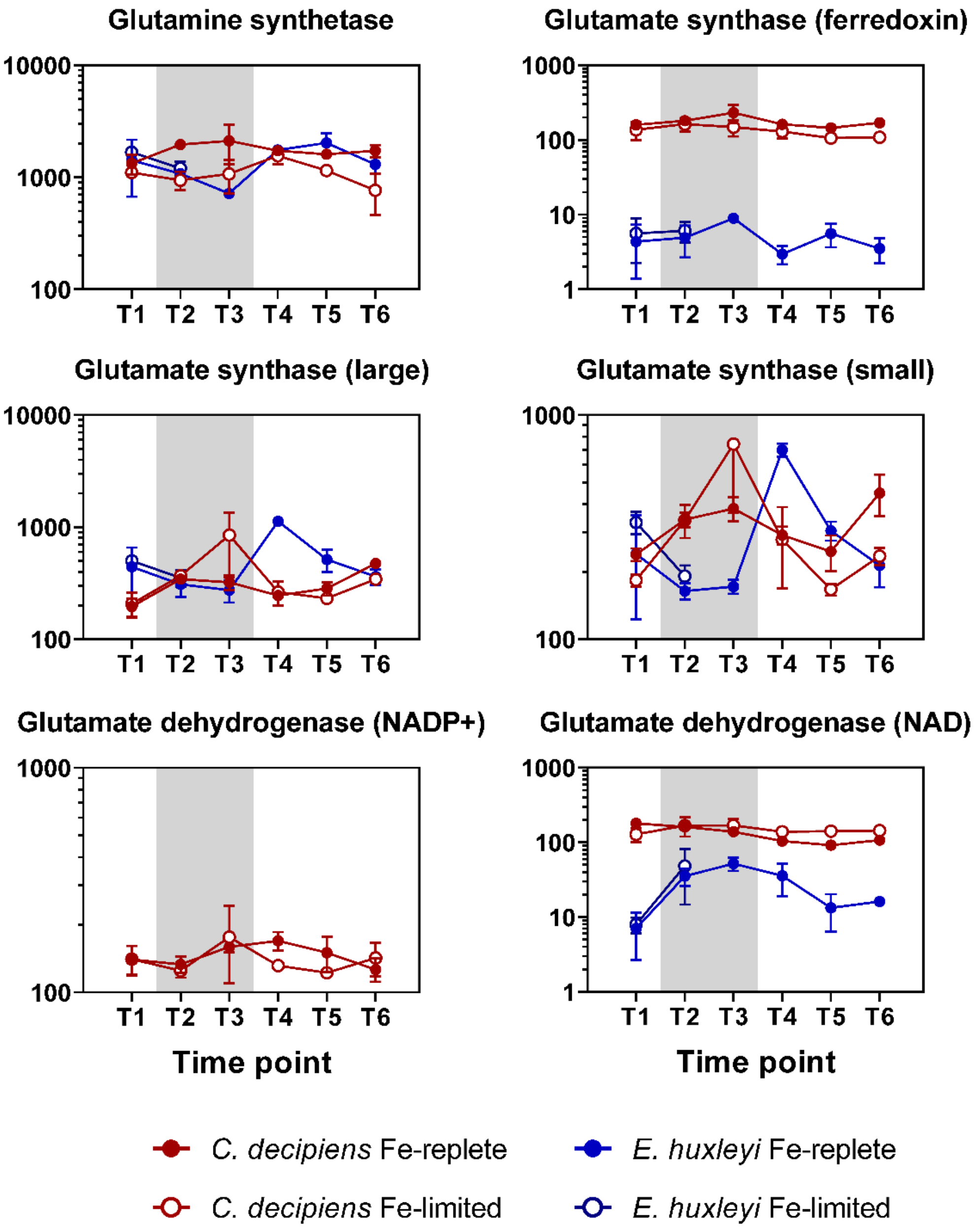

**Fig. S9.** Line graph view of urea and ornithine-urea cycle DESeq2-normalized gene counts in both *C. decipiens* (red) and *E. huxleyi* (blue) under Fe-replete (closed circles) and Fe-limited (open circles) conditions. Gray background shading indicates measurements for time points in the dark. Error bars indicate the standard deviation of the mean where n = 3 except *C. decipiens* iron-replete T5 and *E. huxleyi* Fe-replete T3 where n = 2.

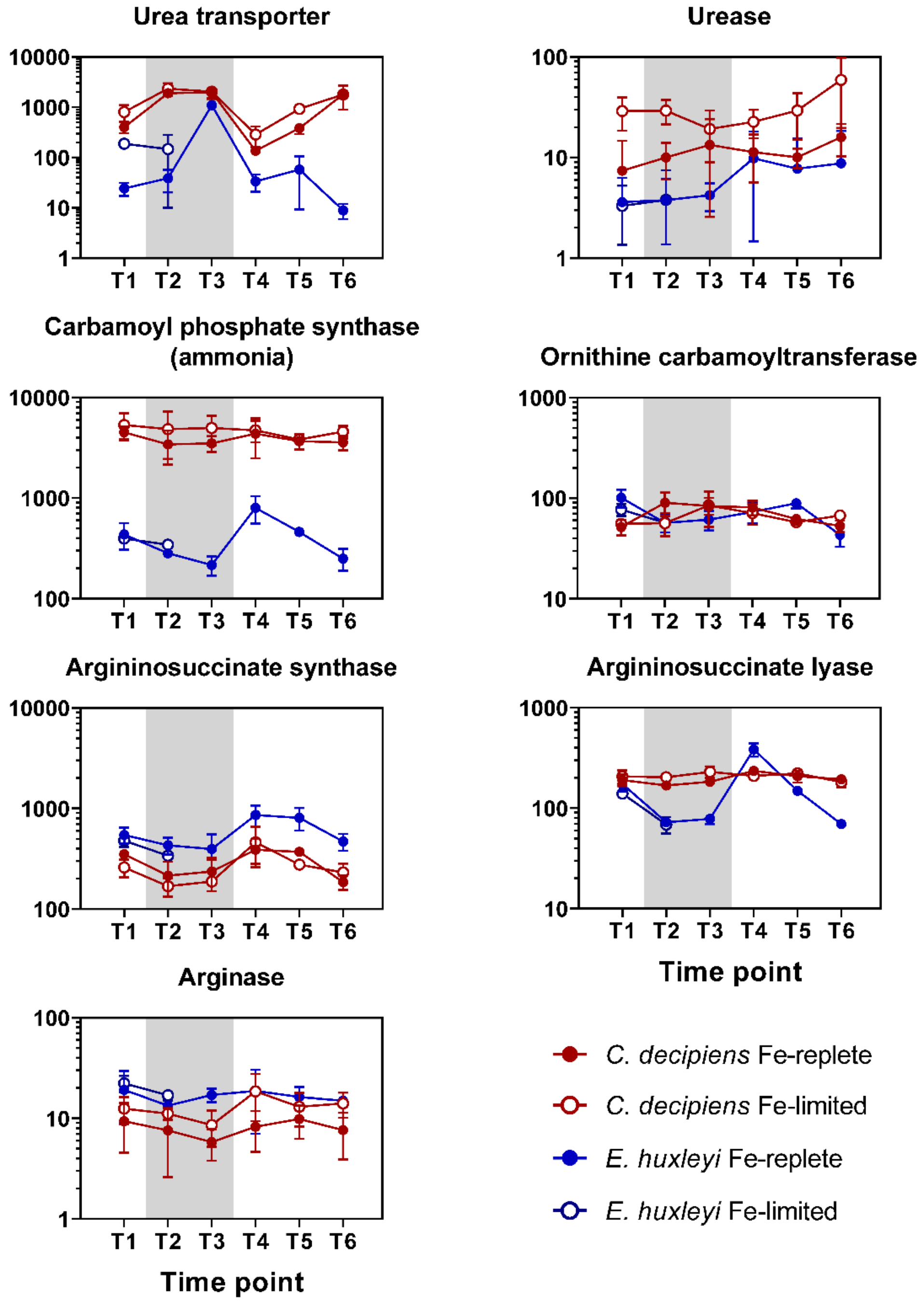

**Fig. S10.** Differential expression for nitrogen-related genes between iron treatments (Fe-replete / Fe-limited) at each time point for *C. decipiens* and *E. huxleyi*. Only T1 and T2 were analyzed in *E. huxleyi* as that is where triplicate samples were available. Significance in differential expression is shown within each cell as follows: *P ≤ 0.05, **P ≤ 0.01, ***P ≤ 0.001, ****P ≤ 0.0001.

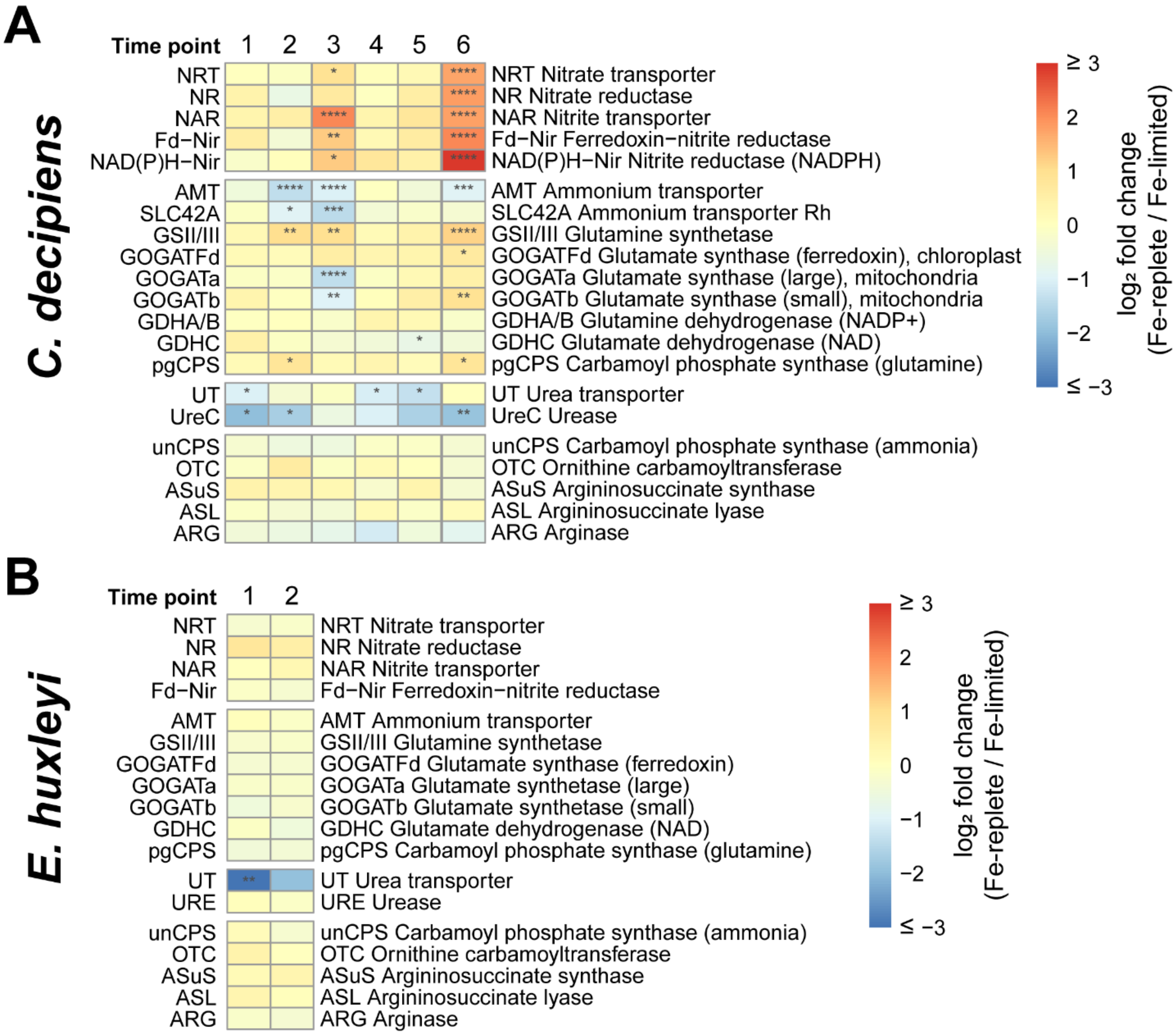

**Fig. S11.** Scale-free fit index as a function of the soft-thresholding power calculated with the pickSoftThreshold function in WGNCA (1). The red horizontal line indicates the cutoff for selecting the soft-thresholding power ($R^2 = 0.9$).

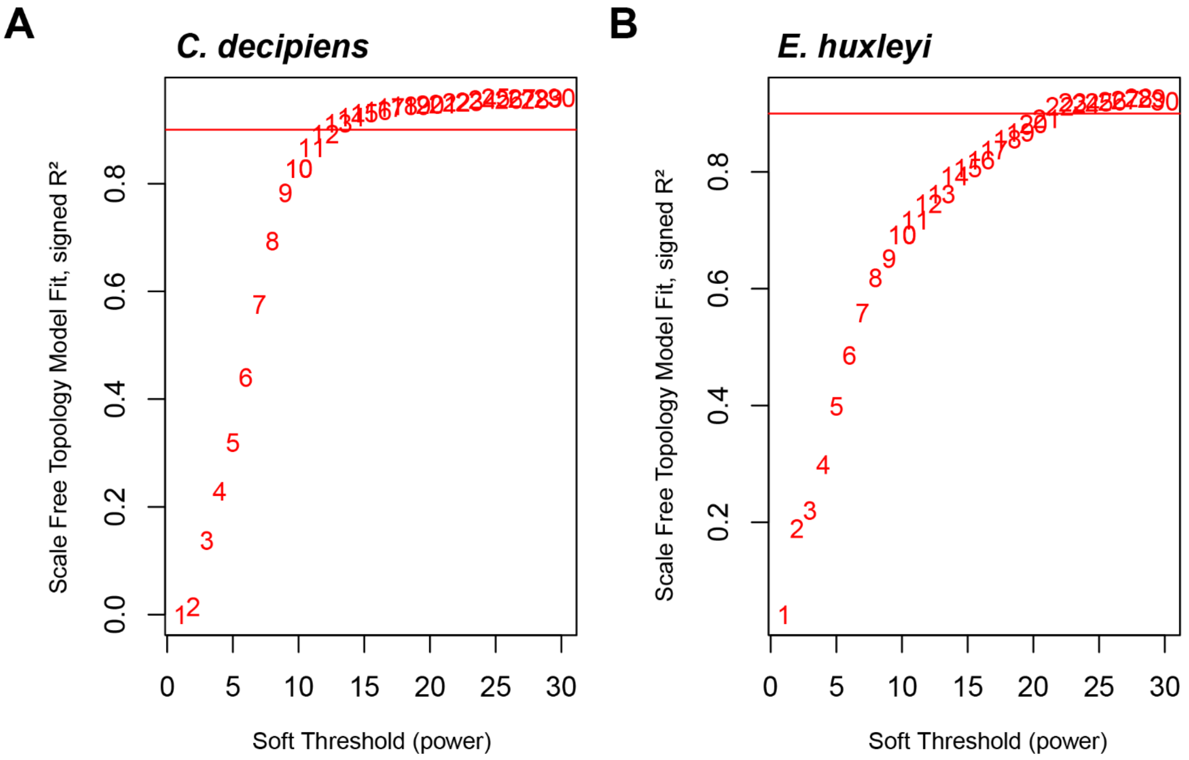

**Table S1.** Reference transcriptome sequencing statistics for each Illumina run on both isolates.

| Organism | Raw Reads | Trimmed Reads | Average trimmed read length (bp) |
|---|---|---|---|
| ***Chaetoceros decipiens*** | | | |
| HiSeq (High Output) | 15,106,132 | 13,944,800 | 99 |
| HiSeq (Rapid Run) | 7,604,228 | 4,619,784 | 150 |
| MiSeq | 1,722,190 | 183,444 | 232 |
| **Total** | **24,432,550** | **18,748,028** | |
| ***Emiliania Huxleyi*** | | | |
| HiSeq (High Output) | 20,890,580 | 19,027,108 | 99 |
| HiSeq (Rapid Run) | 11,865,354 | 7,820,588 | 148 |
| MiSeq | 1,811,203 | 196,508 | 168 |
| **Total** | **34,567,137** | **27,044,204** | |

**Table S2.** Reference transcriptome assembly statistics for both isolates. Contigs, isoforms, and contig N50 are based on the initial assembly. The number of contigs clustered by 99% similarity and predicted proteins from the clustered contigs is also shown. The number and percentage of predicted proteins annotated for each protein database are provided. BUSCO scores were calculated with BUSCO v4.1.4 using the Eukaryota, Stramenopile, and Chlorophyta datasets (2).

| | ***Chaetoceros decipiens* UNC1416** | | ***Emiliania huxleyi* UNC1419** | |
|---|---|---|---|---|
| **Contigs** | 53,420 | | 112,391 | |
| **Isoforms** | 48,500 | | 81,159 | |
| **Contig N50** | 343 | | 975 | |
| **Contigs (99% similarity)** | 50,817 | | 98,927 | |
| **Predicted proteins** | 11,667 | | 36,968 | |
| **Predicted protein N50** | 546 | | 957 | |
| **Predicted proteins with annotations** | | | | |
| KEGG | | | | |
| *Genes* | 8,241 | 70.6% | 30,585 | 82.7% |
| *Orthologs* | 4,431 | 38.0% | 12,664 | 34.3% |
| *Unique Orthologs* | 2,437 | | 4,169 | |
| *Modules* | 1,371 | 11.8% | 2,461 | 6.7% |
| UniProt | 8,505 | 72.9% | 31,092 | 84.1% |
| Pfam | 4,960 | 42.5% | 16,796 | 45.4% |
| Unique Pfam | 2,019 | | 3,288 | |
| PhyloDB | 8,269 | 70.9% | 31,439 | 85.0% |
| **BUSCO** | | | | |
| Eukaryota | | | | |
| *Complete* | 51 | 20.0% | 167 | 65.5% |
| *Fragmented* | 73 | 28.6% | 37 | 14.5% |
| *Total* | 124 | 48.6% | 204 | 80.0% |
| Stramenopile | | | | |
| *Complete* | 41 | 41.0% | 85 | 85.0% |
| *Fragmented* | 24 | 24.0% | 8 | 8.0% |
| *Total* | 65 | 65.0% | 93 | 93.0% |
| Chlorophyta | | | | |
| *Complete* | 391 | 25.7% | 985 | 64.8% |
| *Fragmented* | 101 | 6.6% | 45 | 3.0% |
| *Total* | 492 | 32.4% | 1,030 | 67.8% |
| **Predicted proteins with reads** | | | | |
| Reference reads | 11,662 | 100.0% | 36,496 | 98.7% |
| Tag-Seq reads | 11,648 | 99.8% | 35,494 | 96.0% |

**Table S3.** Software version numbers and parameters used. If not stated, the default options were used.

| **Software** | **Version** | **Options** |
|---|---|---|
| Trimmomatic (reference transcriptome) | 0.36 | ILLUMINACLIP:TruSeq3-PE.fa:2:30:10<br>LEADING:3<br>TRAILING:3<br>SLIDINGWINDOW:4:15<br>MINLEN:36 |
| Trimmomatic (TagSeq reads) | 0.38 | ILLUMINACLIP:TruSeq3-PE.fa:2:30:10<br>LEADING:3<br>TRAILING:3<br>SLIDINGWINDOW:4:15<br>MINLEN:20 |
| Trinity | 2.5.1 | --SS_lib_type FR<br>--min_contig_lenth 90 |
| CD-HIT-EST | 4.7 | -c 0.99 |
| Genemark S-T | | --fnn<br>--faa |
| BLASTP | 2.7.1 | -max_target_seqs 10<br>-evalue 0.00001 |
| hmmscan | 3.1b2 | --cut_ga |
| Salmon index | 0.9.1 | --type quasi<br>-k 15 |
| Salmon quant | 0.9.1 | -l U<br>--gcBias |

**Dataset S1.** Sequencing results for each experimental sample with corresponding NCBI SRA information.

**Dataset S2.** WGCNA results for *C. decipiens*. Module membership is defined as the Pearson correlation between the contig and the module eigengene. Gene TP significance is defined as the Pearson correlation of between the contig and the time point. P-values were corrected for multiple testing using the Benjamini & Hochberg false discovery rate controlling procedure (3).

**Dataset S3.** WGCNA results for *E. huxleyi*. Module membership is defined as the Pearson correlation between the contig and the module eigengene. Gene TP significance is defined as the Pearson correlation of between the contig and the time point. Corresponding p-values were corrected for multiple testing using the Benjamini & Hochberg false discovery rate controlling procedure (3).

**Dataset S4.** Results from ANOVAs for data displayed in Figure 2.

**Dataset S5.** Differential expression results between iron treatments (iron-replete / iron-limited) for *C. decipiens.*

**Dataset S6**. Differential expression results between iron treatments (iron-replete / iron-limited) for *E. huxleyi*.
